# Dynamic organization of cerebellar climbing fiber response and synchrony in multiple functional modules reduces dimensions for reinforcement learning

**DOI:** 10.1101/2022.12.05.518634

**Authors:** Huu Hoang, Shinichiro Tsutsumi, Masanori Matsuzaki, Masanobu Kano, Mitsuo Kawato, Kazuo Kitamura, Keisuke Toyama

**Author notes:** Correspondence: Kazuo Kitamura and Keisuke Toyama. These authors contributed equally to this work (Huu Hoang, Shinichiro Tsutsumi).

## Abstract

Daynamic functional organization by synchronization is theorized to be essential for dimension reduction of the cerebellar learning space. We analyzed a large amount of coordinate-localized, two-photon imaging data from cerebellar Crus II in mice undergoing “Go/No-go” reinforcement learning. Tensor component analysis revealed that a majority of climbing fiber inputs to Purkinje cells were reduced to only four functional components, corresponding to accurate timing control of motor initiation related to a Go cue, cognitive error-based learning, reward processing, and inhibition of erroneous behaviors after a No-go cue. Spatial distribution of these components coincided well with the boundaries of Aldolase-C/zebrin II expression in Purkinje cells, whereas several components are mixed in single neurons. Synchronization within individual components was bidirectionally regulated according to specific task contexts and learning stages. These findings suggest that the cerebellum, based on anatomical compartments, reduces dimensions by self-organization of components, a feature that may inspire new-generation AI designs.

## Introduction

Computational learning theory asserts that machine learning algorithms necessitate as many training data samples as the number of their parameters for correct generalization^1–5^. In contrast, although the cerebellum contains tens of billions of neurons and even more synapses, and is involved in diverse functions^6, 7^, each of which may require a different coding scheme^8–10^, animals can learn new behaviors within thousands of trials, for which the cerebellum is mainly responsible^11^. To reconcile these observations, previous theories have proposed that cerebellar compartments^12–14^ and spike synchronization drastically reduce effective numbers of learning parameters (degrees of freedom) and enable learning from small samples^15–18^. On one hand, the cerebellar cortex is organized into multiple longitudinal compartments with different intrinsic neuronal activity^12, 19–25^, each forming a computational unit^26^. Outstandingly in each of these, aldolase C (AldC)-positive and negative zones represent distinct information in adaptation of the vestibulo-ocular reflex^27–29^ and in eyeblink conditioning^30–32^. Therefore, this compartmentalization constitutes, at least in part, dimension reduction for cerebellar functions related to eye movement and conditioning. On the other hand, synchronization of climbing fiber (CF) activity, which mainly originates from inferior olivary neurons^33–35^, contributes strongly to cerebellar functions^36–38^. During acquisition of complex behaviors in motor activity^39–41^ or reward processing^42–45^, CF synchrony could provide flexible dimension control of neural dynamics^15, 16, 18, 46^. Moreover, synchrony is high within cerebellar compartments, but not across them, implying a structure-function relationship between cerebellar compartments and synchronization^47–49^. However, it is unclear how compartments and synchronization together might facilitate dimension reduction for learning of cognitive functions for which genetic prewiring is unlikely to be sufficient.

To shed light on mechanisms of dimension reduction in the cerebellum, we systematically examined two-photon recordings of CF activity from Purkinje cells in eight AldC compartments of mice learning an auditory discrimination Go/No-go task^50^. This task requires reinforcement learning guided by reward, and necessitated, first, accurate timing control of licking to obtain a reward, second, learning based on reward prediction error, third, reward processing, and fourth, inhibition of licking to the No-go-cue. We applied a hyper-acuity spike timing detection algorithm with 10-ms resolution^51^ and tensor component analysis^52^ of CF firings, and found that 50% of the variance was explained by only four clearly separable components, each corresponding to one of the above four functions. Furthermore, spatial distributions of these four components were markedly different with functional boundaries between the medial and lateral Crus II and across AldC compartments. Ten-millisecond (ms) resolution analyses revealed that the CF synchrony of a specific component was high in a number of synchronous neurons and in synchrony strength for a specific engaged cue-response condition, and was spatially localized within corresponding compartments. Strikingly, tensor components were shaped by bidirectional synchrony-response changes during the course of learning. These results demonstrated that the cerebellum reduces dimensions of activity in a large number of neurons to a much smaller number of components, each of which is driven by a synchronization scheme that conforms to a specific task. Interestingly, we also found that individual anatomical zones and even a single CF could contain signals from multiple components^53–56^.

This study provided the first evidence simultaneously supporting the two major theories of cerebellar functions in a single task^18, 57–62^, and should contribute to resolution of the long-standing controversy^63^. This study also unveiled the secret of cerebellar functional architecture: learning from small samples is achieved by compartmentalization (reduced degrees of freedom) due to synchronization and dynamics; therefore, it may contribute to new-generation AI designs.

## Results

### Two behavioral learnings in a Go/No-go task

We trained mice to perform a Go/No-go auditory discrimination task (Figure 1A), which required cognition beyond sensorimotor-related functions, thereby revealing functional differences across AldC compartments during learning^50^. Briefly, mice (n = 17) were trained to associate the “Go” cue (a 9-kHz tone for 0.5 s) with a water reward and to react by licking during a response period of 1 s after cue onset. The “No-go” cue (a 4-kHz tone for 0.5 s) was not associated with a reward, but a 4.5-second timeout was imposed if the mice licked during the response period (Figure 1A). We recorded 87 sessions from 17 mice (each underwent no more than 7 sessions), including 26,517 trials of auditory discrimination Go/No-go tasks in the three learning stages (1st, 2nd, and 3rd stages with fraction correct <0.6, 0.6 - 0.8, 0.8<,respectively). We categorized trials into the four cue-response conditions: HIT trials in the go task (lick after go cue, n = 12,334), false alarm (FA) trials (lick after No-go cue, n = 5,588), correct rejection (CR) trials (no lick after No-go cue, n = 7,681) and MISS trials (no lick after go cue, n = 914). Note that before the main Go/No-go auditory discrimination training, mice had already been trained to lick in response to both high and low tones with indiscriminate reward feedback.

**Figure 1:**
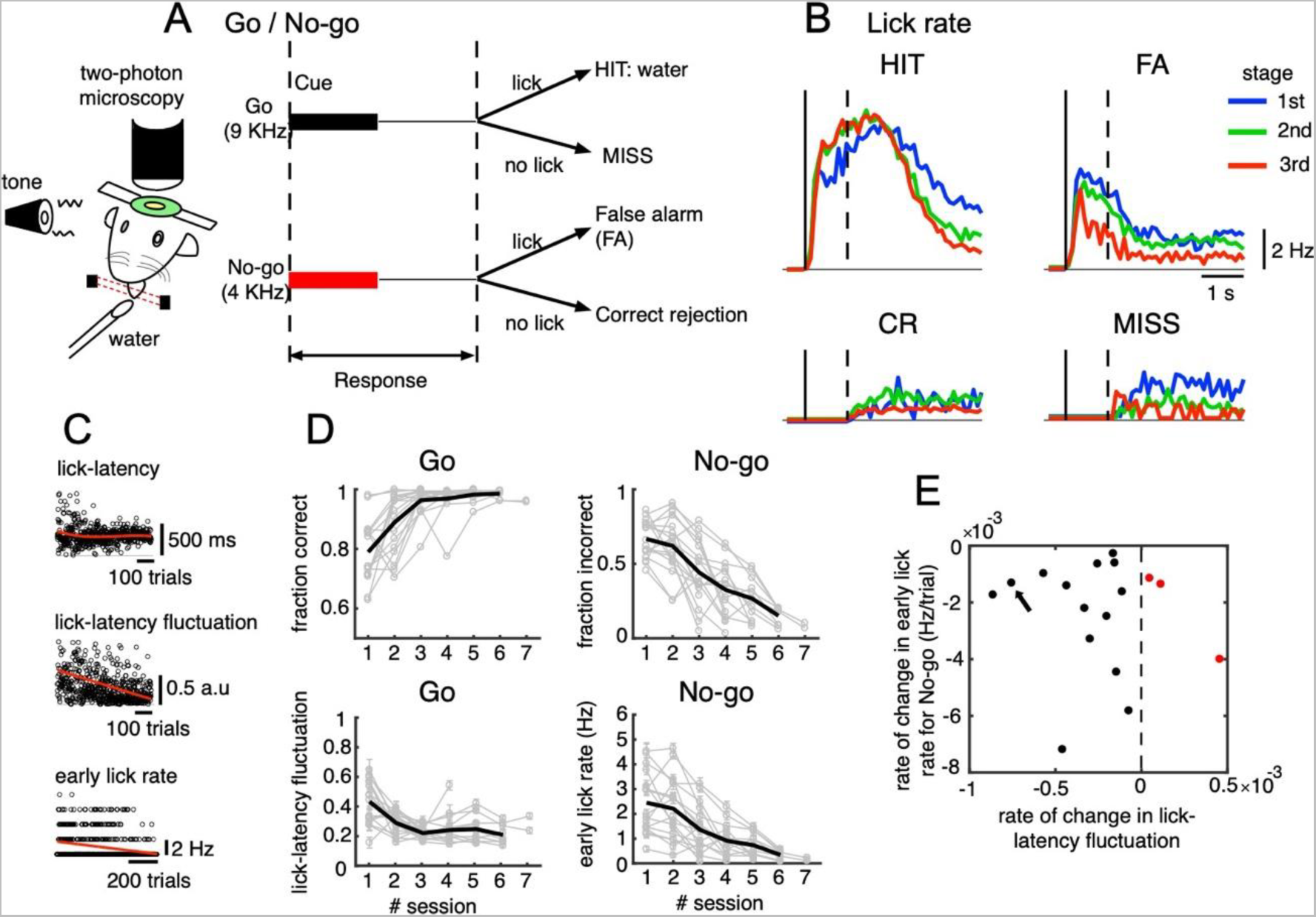
Go/No-go auditory-cue discrimination task and behavior changes during learning. A: Schematic diagram of a mouse performing the Go/No-go discrimination task under a two-photon microscope. B: the lick rate of the four cue-response conditions sampled in the three learning stages (blue, green and red traces for 1st, 2nd and 3rd stages, respectively). C: from top to bottom, lick-latency, lick-latency fluctuation in Go trials and the early lick rate in No-go trials of a representative mouse (indicated by black arrow in E). Trials were sorted by the time course of training. Red traces indicate polynomial fittings of lick parameters as functions of trials (see Methods). D: changes in four learning indices as functions of training sessions, including the fraction correct of Go cues, the fraction incorrect of No-go cues, lick-latency fluctuation in Go trials, the early lick rate in No-go trials. Thin gray traces are of individual animals (n = 17 mice) and thick dark traces are the means. E: scatter plot for rate of change in lick-latency fluctuation for Go cues (abscissa) and rate of change in early lick rate for No-go cues (ordinate) estimated from licking behavior of individual mice. Black dots were for mice whose rates were both negative and red dots were for the three mice that showed increased lick-latency fluctuation after learning (positive rate).

Behavioral data indicated that the lick rate was dramatically reduced in FA trials during the three stages of learning (Fig 1B), but it remained relatively unchanged in HIT trials. During learning, there was an increase in the fraction correct for the Go cue (0.78 ± 0.1 and 0.98 ± 0.01, 1st and 6th sessions, respectively, Fig 1D). There were no consistent changes in the reaction time after Go; however, we found decreased lick-latency fluctuation in the licking response with learning (0.45 ± 0.15 and 0.22 ± 0.06, 1st and 6th sessions, respectively, Fig 1C-D). For the No-go cue, the fraction incorrect decreased after learning (0.66 ± 0.12 and 0.15 ± 0.06, 1st and 6th sessions, respectively, Fig 1D), along with a decreased lick rate in the early response window (0-500 ms after cue onset, 2.4 ± 1.2 and 0.3 ± 0.2 Hz, 1st and 6th sessions, respectively, Fig 1C-D). At an individual level (see Fig S1 for behavior changes associated with Go and No-go cues of individual mice), 14 out of 17 mice showed a negative rate of change in lick-latency fluctuation, while the rate of learning-related changes in the number of early licks for the No-go cue was negative for all mice (Fig 1E). In summary, behavioral data indicated that mice successfully learned an auditory discrimination task by changing licking behavior in the early response window: first, achieving more precise timing of the first lick after Go cues and second, suppressing licks after No-go cues.

### Opposite changes in cue-related CF responses in medial and lateral parts of Crus II during learning

We used the Aldoc-tdTomato transgenic mouse line^49^ to systematically explore functional differences in CF inputs to AldC compartments at single-cell resolution during the task. While mice underwent the Go/No-go task, we performed a total of 236 sessions of two-photon calcium imaging (sampling rate, 7.8 Hz) from PC dendrites at every boundary of AldC expression in eight AldC compartments (7+, 6-, 6+, 5-, 5+, 5a-, 5a+ and 4b-) in lobule Crus II to simultaneously monitor CF-dependent dendritic Ca^2+^ signals (see Methods and Tables S1-2 for detailed numbers of neurons and trials recorded). To investigate CF signals with higher temporal resolution than the two-photon recordings, CF firings were estimated for 6,445 PCs using hyperacuity software^51^ (HA_time, 100 Hz, see Methods and Fig S2A for reliability of HA_time). These technical advances allowed us to monitor CF firing activities of a large number of Purkinje cells in different AldC compartments during the course of learning with high temporal precision.

We studied population peri-stimulus time histograms (PSTHs) of CF firings sampled in the three learning stages for the four cue-response conditions. We found contrastive response profiles between the lateral (AldC 7+ to 5-) and medial parts of Crus II (AldC 5+ to 4b-) segregated by the anatomical and functional border^50^ (Figs 2A-D). CF firings in HIT trials were large and diffusive at the initial learning stage across the entire medial Crus II, as well as a fraction (AldC 6+) of the lateral Crus II (Fig. 2A 1st stage). However, they became even stronger and compartmentally more focused on AldC positive compartments of the lateral and medial parts of Crus II (AldC 6+, 5+, 5a+), and temporally confined within 200 ms after cue onset (2nd and 3rd stages). PSTHs in FA trials were as strong as those in HIT trials initially, distributed across almost the entire lateral and medial parts of Crus II, but more in the lateral parts (Fig. 2B, 1st stage). However, they gradually decreased with learning and finally became rather weak (2nd - 3rd stages), while maintaining the initial compartmental distribution profiles. For CR trials, PSTHs were initially localized in the lateral Crus II, and finally became confined to a fraction (AldC 6-, 6+) of the lateral Crus II (Fig. 2C 1st - 3rd stages). There was only spontaneous CF activity in MISS trials across the entire learning stage (Fig 2D).

**Figure 2:**
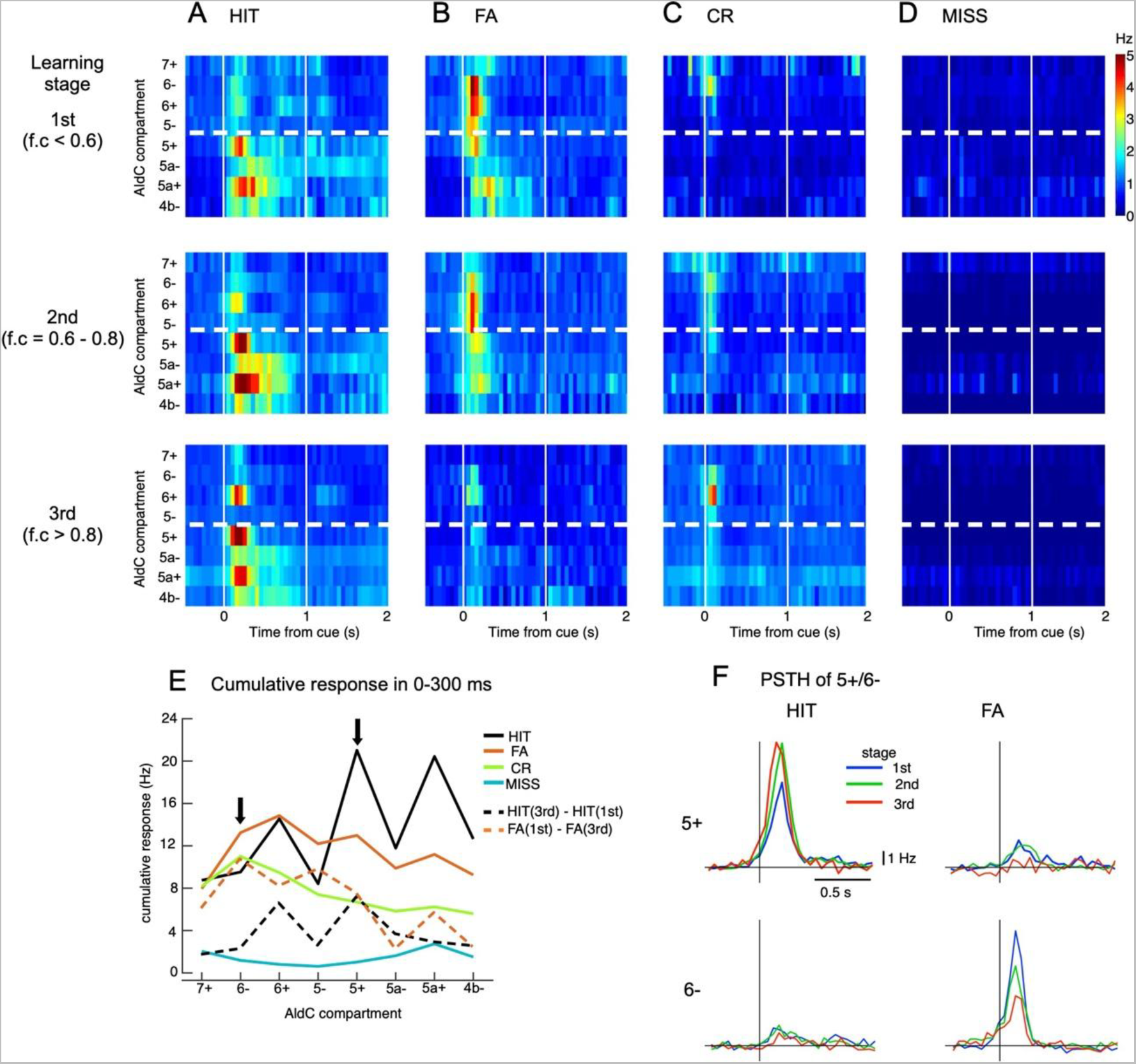
Opposite changes in CF response to cues in the lateral and medial parts of Crus II. A-D: Pseudo-color representation of population peri-stimulus time histograms (PSTHs) of CF responses sampled during three learning stages across four cue-response conditions (HIT trials (A), FA trials (B), CR trials (C), and MISS trials (D)) in each AldC compartment in Crus II. E: response strength estimated as cumulative PSTHs for 0-300 ms after the cue onset for HIT (black trace), FA (orange trace), CR (green trace), and MISS (cyan trace) trials across all three learning stages. Dashed traces show differences in cumulative PSTHs between the 1^st^ and 3^rd^ stages for HIT (black: 3^rd^-1^st^) and FA (orange: 1^st^-3^rd^) for individual AldC compartments. Black arrows indicate the two representative AldC compartments that show the largest changes in CF response after learning for HIT (5+) and FA (6-) trials. F: PSTHs in HIT and FA trials of the two representative AldC compartments 5+ and 6- during learning.

We also studied the compartmental topology of the cue-response for the population PSTH, including all three learning stages, evaluating response strength as cumulative PSTHs for 0-300 ms after cue onset (Fig. 2E, see Methods). Response strength in HIT trials (black trace) across compartments exhibited 3 high peaks – including compartments 5+ (21 Hz), 5a+ (20.5 Hz), and6+ (14.4 Hz) – and 4 smaller valleys – including compartments 7+ (8.6 Hz), 5- (8.4 Hz), 5a- (11.8 Hz), and 4b- (12.4 Hz). By contrast, response strength in FA trials was stronger in the lateral than the medial Crus II. Response strength in CR shows a decline from lateral to medial, and MISS trials remained almost flat across both medial and lateral parts of Crus II.

To better demonstrate opposite response changes of medial and lateral parts at each learning stage, we selected the top 100 neurons in AldC compartments 5+ and 6- (Fig 2E), which showed the largest differences in response strength to HIT and FA trials, respectively, i.e., the neurons whose response strength was most selective for HIT or FA trials (see Methods). We observed that AldC 5+ neurons exhibited a marked increase in PSTHs as well as a phase advance in cue-response along with learning for the HIT condition (peak PSTH, 6.6 and 9.4 Hz for the 1st and 3rd stages, respectively, Figure 2F), while that of AldC 6- neurons showed a significant decrease for the FA condition along with learning (8.6 and 4.0 Hz for the 1st and 3rd stages, respectively). Importantly, we found remarkably opposite changes in synchrony with learning of neurons in AldC compartments 5+ and 6-, which were strongly associated with corresponding opposite changes in their responses even on a single-trial basis (Fig S3). We name them “bidirectional synchrony-response changes”.

### Tensor component analysis of CF activities

We examined whether bidirectional synchrony-response changes observed in AldC compartments 5+ and 6- can be generalized across the entire Crus II. For this purpose, we conducted tensor component analysis^52^ (TCA) to decompose high-dimensional CF firings, i.e., PSTHs of neurons in the four cue-response conditions sampled during three different learning stages, into low-dimensional components, each with a unique set of coefficients of a temporal factor, the cue-response condition, and the neuron (see Methods for more details).

We found that only four distinct tensor components underlie CF firings (Fig 3), and they accounted for about half the variance in 6,445 neuronal PSTHs (see Fig S4 for more details). The first tensor component (TC1) had a fast temporal profile peaking at 200 ms after cue onset (Fig 3A). This component TC1 was dominant mostly in the HIT condition and only weakly manifested in FA (Fig 3B). Compartmentally, TC1 was concentrated in AldC positive compartments across the medial and lateral Crus II and gradually increased its coefficients along with learning (Fig 3C). By contrast, the second tensor component (TC2) was dominant in the FA condition and weakly manifested in HIT (Fig 3A). It had a very fast temporal profile that peaked at 100 ms after cue onset, and decayed sharply toward baseline within 400 ms after cue onset. TC2 was distributed broadly in the lateral Crus II during early stages of learning (Fig 3C), but its coefficients were markedly reduced in the 3rd stage. The third tensor component (TC3) had a slow temporal profile, peaking at ∼300 ms, and was prolonged for 1 s after cue onset. TC3 was dominant in the HIT condition and little observed in FA. Compartmentally, TC3 was initially distributed in the entire medial Crus II, but mainly in AldC negative compartments during later stages of learning (Fig 3C). Finally, the fourth tensor component (TC4) had temporal profile and compartmental distribution similar to those of TC2, but was present only in the CR condition. In summary, TCA revealed four tensor components with distinct temporal and compartmental profiles for different cue-response conditions.

**Figure 3:**
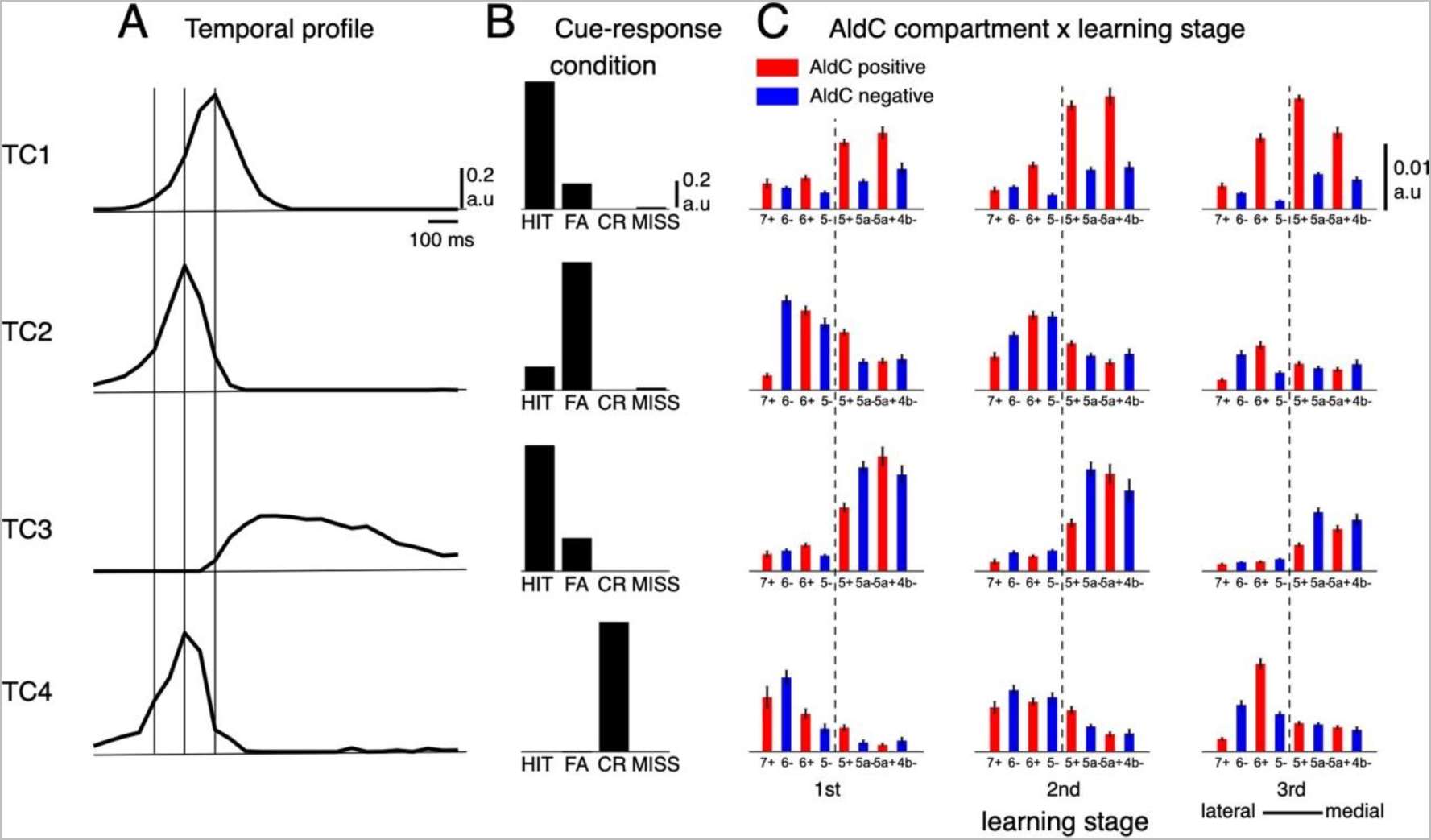
Tensor component analysis of CF activity. A-C: (A) Coefficients of temporal, (B) cue-response condition, and (C) neuronal factor of the four tensor components (TC1-TC4, from top to bottom) estimated by TCA. TCs are shown in their contribution order (see Methods for details). The three vertical lines in (A) represent the timing at 0, 100 and 200 ms after the cue onset. Bars with lines in (C) show means and SDs of neuron factor coefficients, grouped by eight AldC compartments (shown in red and blue colors for AldC positive and negative compartments, respectively) and three learning stages. The thick dashed line in (C) indicates a functional boundary between the lateral and medial Crus II^48^.

### Synchrony dynamics shape tensor component activities

To investigate synchrony dynamics of the four tensor components, we selected the neurons that were most strongly represented by each component, i.e., with the highest contribution by that component. Briefly, we sampled the top 300 neurons for each TC at each learning stage, while rejecting those that overlapped. As a result, we selected 2,096 neurons, whose cumulative responses accounted for approximately 40% of total responses in the entire population (see Methods and Fig S5 for more details; Basically, the same results were obtained when we selected fixed proportion of the sampled neurons for each TC at each learning stage). Firings for each class of neurons varied considerably with learning stages, while maintaining roughly the same temporal profiles of the corresponding tensor components (compare Fig. S5C PSTH temporal profiles at the three learning stages with the corresponding temporal profiles of TCs in Fig 3A).

We found that synchrony strengths (estimated as the summation of cross-correlograms in ±10 ms around the center time bin, see Methods) within TCs (0.40 ± 0.17, 0.36 ± 0.17, 0.33 ± 0.17 and 0.19 ± 0.13 for TC1-4, respectively) were significantly larger than those across TCs (0.12 ± 0.09, p < 0.0001 for all pairwise comparisons between within-TC and across-TC populations, Fig 4A). We further investigated synchrony strengths for specific cue-response conditions in which TC1-2 neurons were engaged. More specifically, synchrony strength (which was normalized by the number of spikes) of TC1 in HIT trials (0.40 ± 0.17) and that of TC2 in FA trials (0.37 ± 0.17) were significantly stronger than those of TC1-2 in other cue-response conditions (0.19 ± 0.11 and 0.23 ± 0.14 for TC1-others and TC2-others, respectively, Fig 4B). That tendency was observed even at a single-trial basis, with strong instantaneous synchrony (total number of synchronous firings in 30-ms bins in a window of 300 ms before the first lick onset, see Methods) was found in TC1-HIT and TC2-FA trials at the 3rd and 1st learning stages, respectively (Fig 4C and Supplemental Movie M1). Furthermore, such synchrony dynamics were well aligned with the spatial distribution of TC1-2 (see Supplemental Movie M2). Together, these results suggested that synchronization, spatially guided by AldC compartments, organizes TC populations only during dedicated cue-response conditions.

**Figure 4:**
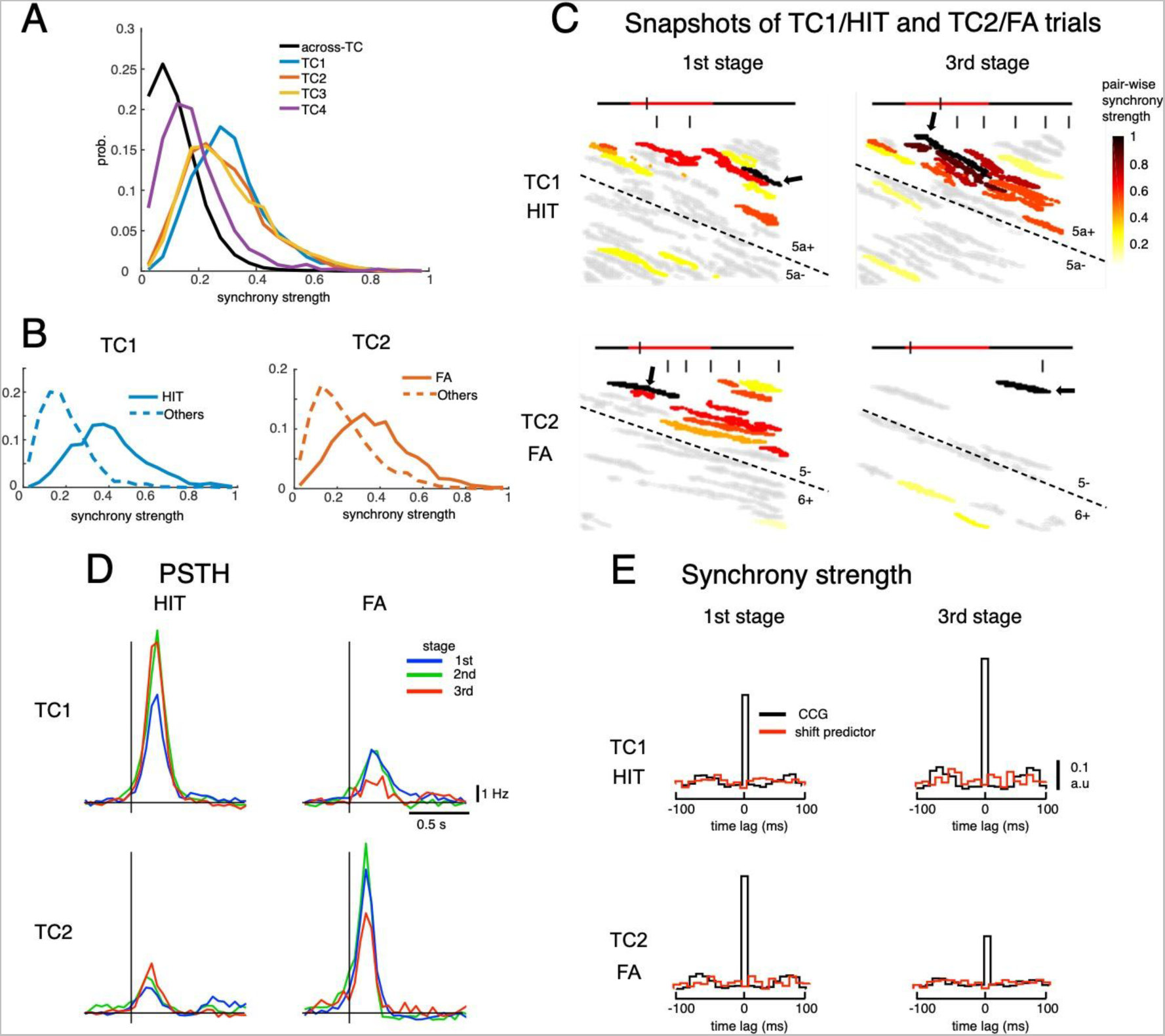
Dynamics of synchrony and opposite changes in synchrony of TC1-TC2 during the course of learning. A: histograms of synchrony strength between TC1-4 neurons (colored traces) in comparison with those across TCs (black trace). B: histograms of synchrony strength computed specifically in HIT trials for TC1 (solid blue trace) and in FA trials for TC2 (solid orange trace), contrasting with those in other cue-response conditions (dashed traces). C: representative images of synchronous firings in TC1/HIT (5a−/5a+) and TC2/FA (6+/5−) in the 1st and 3rd stages. The horizontal line shows the time course of −200 ms to 1 s after cue onset with a small tick indicating the timing of the snapshot relative to the cue onset (the cue period of 500 ms was shown in red color). Short vertical bars indicate the lick timings. The snapshots capture firings of Purkinje cell dendrites (gray areas) co-activated in a time bin of 10 ms. The hot-color spectrum represents pair-wise synchrony strength between the reference cells (pointed by black arrows) and other cells in the same recording session (see Supplemental Movies). D: PSTHs of TC1-TC2 neurons in HIT and FA trials in three learning stages. E: population CCGs of the 1^st^ and 3^rd^ stages indicated opposite changes in synchrony strength of TC1-TC2 neurons during learning. The red traces in CCGs indicate shift predictors estimated for the correlation solely due to the cue stimulus.

Moreover, we found that the firing of TC1 strongly increased (peak PSTH, 6.8 and 10.5 Hz for 1st and 3rd stages, respectively, Fig 4D) along with learning for the HIT condition, while it was moderate at the 1st stage and almost disappeared at the 3rd stage for the FA condition. By contrast, the cue-response of TC2 remained small across all learning stages for the HIT condition, while it was strong at the 1st and 2nd stages and markedly decreased at the 3rd stage for the FA condition (10.6 and 6.8 Hz for 1st and 3rd stages, respectively, Fig 4D). We also observed clear synchrony changes during learning only for neurons of TC1 (increased for Go trials) and TC2 (decreased for No-go trials, Fig 4E and S6). Together, those findings suggest that synchronization may drive two opposite response changes in the early window associated with the Go cue for TC1 and the No-go cue for TC2.

### Correlations between CF synchrony and licking behaviors

The facts that TC1 increased synchrony in HIT trials during learning of precisely timed initiation of licking and that TC2 decreased synchrony in FA trials along with reduction in erroneous licks with learning suggest that two opposite synchrony changes are related to two changes in licking behaviors for the two corresponding cue-response conditions. To test these possibilities, we investigated synchrony-behavior correlations in the three response windows, namely, early lick (0 – 0.5 s), reward lick (0.5 – 2 s) and succeeding lick (2 – 4 s). These windows were determined from behavioral data showing that rewards were given in a window of 0.41-1.23 second after the first lick, timing of which was about 0.5 s after cue onset.

For the early lick window, we found a tendency for TC1-synchronous spike-triggered lick responses in Go trials to be mostly positive, peaking at 200-300 ms for all three learning stages (Fig 5A). Importantly, peak amplitude increased with learning, from 7.4 to 11.5 Hz for the 1st and 3rd stages, respectively. Furthermore, on a single trial basis, instantaneous synchrony of TC1 neurons was negatively correlated with lick-latency fluctuation (slope = -0.02, p < 0.0001, 2,115 trials, see Methods for details).

**Figure 5:**
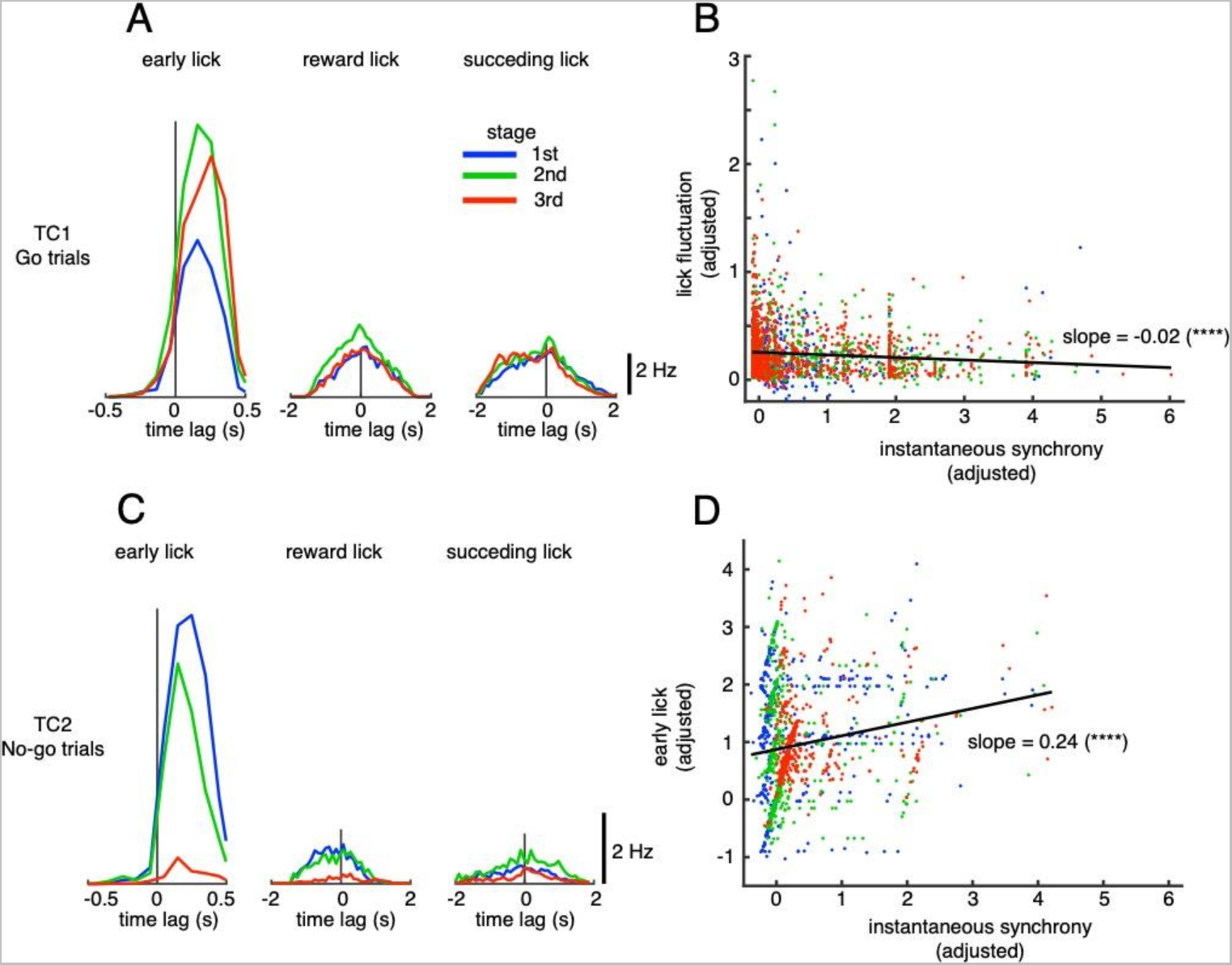
Correlations of TC synchronous activities and licking behavior. A: synchronous spike-lick CCGs of TC1 neurons in the three response windows: early lick (0–0.5 s after cue onset), reward lick (0.5–2 s) and succeeding lick (2–4s). B: scatter plot of instantaneous synchrony and lick-latency fluctuation in Go trials of TC1 neurons. C: synchronous spike-lick CCGs of TC2 neurons in the three response windows similar to those in A. D: scatter plot of instantaneous synchrony and number of early licks in No-go trials of TC2 neurons. Each dot in scatter plots of B-D corresponds to a single trial. We used a multiple linear regression model with the two learning variables as functions of instantaneous synchrony of the four TCs and fraction correct. The black trace represents the correlation of two learning variables and instantaneous synchrony, with a slope and significance level indicated by asterisks. Note that the ordinate and abscissa of scatter plots in A-B were adjusted to show correlations specific to TC1 or TC2 neurons (see Methods).

Conversely, for TC2 neurons in No-go trials, synchronous spike-triggered lick responses in the positive time domain were very strong initially, but disappeared at the later stage (peak amplitude, 7.5 and 0.7 Hz for the 1st and 3nd stages, respectively, Fig 5B). Regression analysis further showed that instantaneous synchrony of TC2 neurons was positively correlated with the early lick rate in No-go trials (slope = 0.24, p < 0.0001, 965 trials Fig 5B). Note that synchronous spike-triggered lick responses of TC1-TC2 in the reward lick and succeeding lick windows were small and unchanged during learning, suggesting that TC1-TC2 were neither related to rewards nor their sensorimotor feedback. Moreover, multiple regression analysis of the two early lick variables (Go and No-go) with instantaneous synchrony of the four TCs combined indicated that lick-latency fluctuation in Go trials was only correlated with synchrony of TC1 neurons, whereas the early lick rate in No-go trials was most strongly correlated with synchrony of TC2 neurons, although moderate correlations with synchrony of TC1 and TC4 neurons were also observed (see Fig S7).

We further conducted a decoding analysis to test how well the spiking model could predict licking behavior. By assuming that a single spike independently triggers a lick event following the spike-triggered lick histogram, we derived the likelihood of a trace of licking events given the spike train (see Supplemental Information for details). These results indicated that synchronous spikes of TC1-2 predicted occurrence of lick events statistically better than all spikes of TC1-2 neurons, all spikes of all neurons in the same recording session, or the chance level for which no correlation between spike and lick events was assumed (Fig S8). At the individual trial level, synchronous spikes of TC1 and TC2 were the best model for approximately 50% of all lick events in HIT and FA trials, respectively (Fig S8C). In addition to the decoding analysis, 4 out of 5 animals showed an increase in timing fluctuation of the first-lick in HIT trials following muscimol injection at the left Crus II (Fig S9). Together, these results suggest strong relationships between synchronization of TC1-2 and licking behaviors.

### Tensor representation in CF responses of individual cells in the cerebellar cortex

While TCA clearly decomposed CF responses into four TCs, we also observed overlaps in TC representation at the individual neuron level, i.e., a single neuron may represent more than one TC by having large coefficients of multiple TCs (see Fig S5A for the ratio of overlapping neurons among TCs). We visualized such overlap by mixing the coefficients of TC1-4 for individual neurons by CMYK colors (Fig 6A). Compartmental distributions of each TC were roughly maintained as shown in Fig 3C. TC1 was mostly distributed in AldC-positive compartments, TC2 and TC4 in the lateral Crus II, and TC3 in the medial Crus II, and mainly in AldC negative compartments at the later learning stage. However, neurons representing different TCs could reside in the same compartments. For instance, TC1 (cyan) and TC2 (magenta) neurons could be found in AldC 6+ in the first two learning stages. More strikingly, we found that a fraction of neurons represented multiple TCs. They were “green” neurons for mixing of TC1-TC3 in AldC compartments 5a- and 5a+, and “dark magenta” neurons for mixing of TC2-TC4 in the lateral Crus II at the first stage. The tensor representation of individual neurons showed that TCs have complicated compartmental distributions that change during learning. At the same time, single AldC compartments and single neurons can represent multiple TCs.

**Figure 6:**
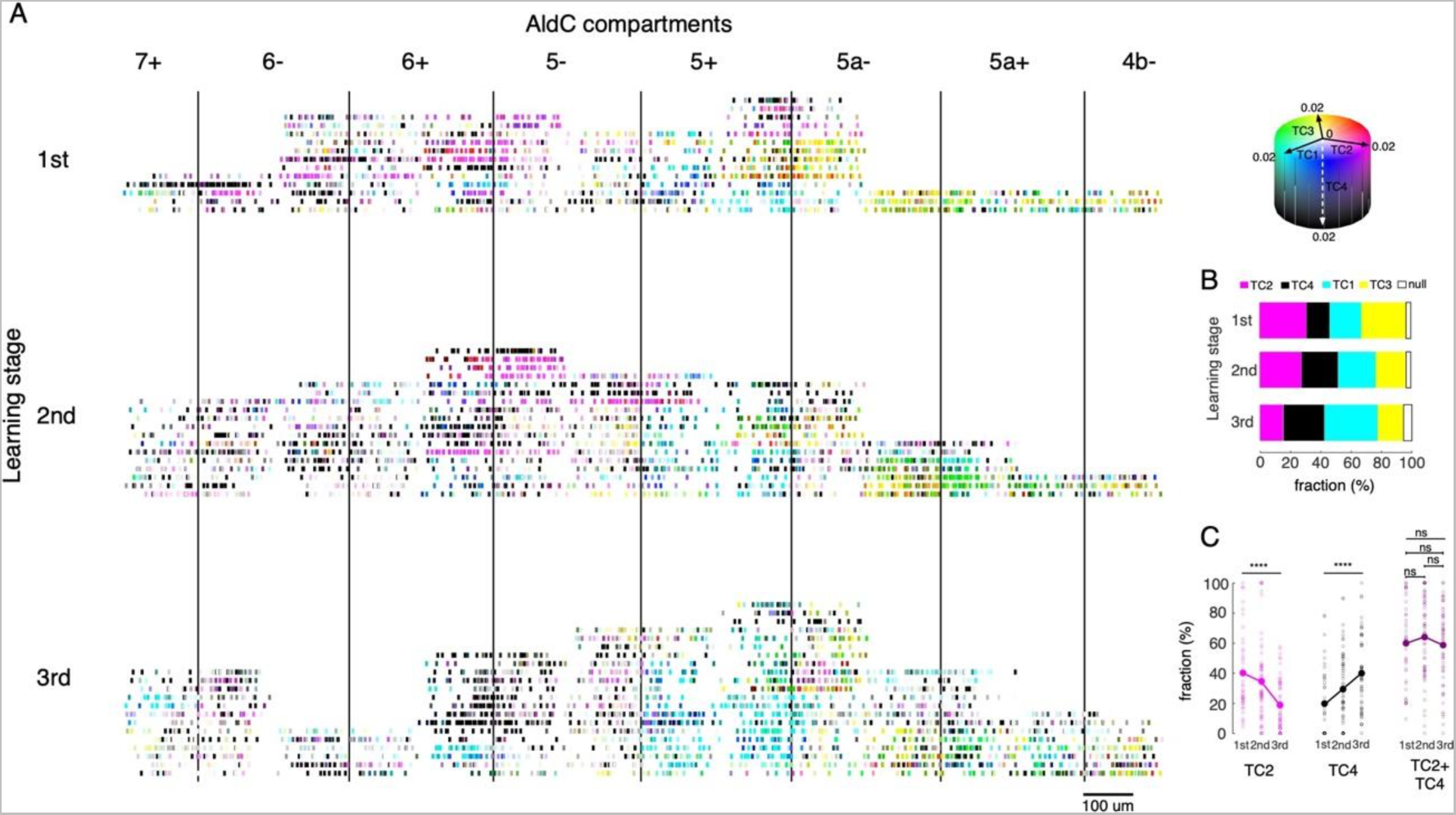
Tensor representation of individual cells in Crus II. A: CF inputs to 6,465 neurons were recorded in 8 AldC compartments (columns) and at three learning stages (rows). Each short bar indicated the location of a single cell relative to the AldC boundaries (vertical lines). The color of each cell was mixed by coefficients of the four tensor components (cyan – TC1, magenta – TC2, yellow – TC3 and black – TC4). Each row corresponds to a single recording session. For visualization purposes, the width of each AldC compartments was manually adjusted to 300 𝜇m. B: fractions of neurons classified as TC1-4 in each of the three learning stages (color bars). Note that less than 6% of all recorded neurons could not be classified as TC1-4 (open bars). C: fractions of neurons in the lateral Crus II (AldC compartments 7+, 6-, 6+, 5-) classified as TC2 (magenta) and TC4 (black) and their summation (TC2 + TC4, dark magenta) in 150 sessions (small open circles) and their means (solid large circles) for the three learning stages.

To systematically investigate changes in TC representation with learning, we classified all recorded neurons into one of the TC1-4 populations based on their TC coefficients, e.g., a particular neuron is classified as TC1 if its coefficient of TC1 is the highest among four TC coefficients. As expected, fractions of TC1 and TC4 neurons increased and those of TC2 and TC3 decreased significantly (Fig 6B). More interestingly, while fractions of TC2 and TC4 neurons in the lateral Crus II (AldC compartments 7+ to 5-) changed in opposite directions (ANOVA, p < 0.0001 for changes in fractions across the three stages, n = 150 sessions, Fig 6C), their summation remained unchanged at about 60% even at the session level (ANOVA and pairwise t-tests, p > 0.3).

## Discussion

One of the technical advances of this study was that we applied the hyperacuity algorithm^51^ to estimate spike timing from Ca signals with 10-ms resolution. This enabled us to study a huge number of neurons (>6,000) with hyperacuity resolution (10 ms) along a whole process of cognitive and motor learning, while spatially guided by Aldolase-C zones. Tensor component analysis (TCA) of this high spatio-temporal CF data revealed distinct components that may be involved in different learning-related functions. In particular, TC1 consisted of neurons in AldC- positive zones with fast responses in the HIT condition (Fig 3A). Importantly, their synchronous spike-triggered lick responses were almost zero for negative time lag and sharply raised in positive time lag (Fig 5A), suggesting that synchronous TC1 spikes control early licks as motor commands. This hypothesis is further supported by our observation that synchronous TC1 spikes best predict early lick timings in HIT among four different models (Fig S8). Furthermore, increased CF synchrony in TC1 was negatively correlated with the decrease of lick-latency fluctuation (Fig 5B). Together, these findings for TC1 are in good agreement with the synchronization and timing control hypothesis of the cerebellum^34, 39, 41, 57, 64^, according to which, more synchronization could stabilize timing of synchronized motor commands by canceling noisy synaptic inputs to the IO, possibly leading to more precise timing control with less fluctuation^65^.

TCA also revealed the component that may be involved in cognitive learning^66, 67^. Namely, the TC2 population, distributed mostly in the lateral Crus II, showed very fast responses in the FA condition (Fig. 3A). As learning proceeded, TC2 synchronous firings in the FA condition decreased dramatically (Fig 4E). Notably, such a decrease in CF firing of TC2 neurons was highly correlated with a decrease in the early lick rate (Fig 5D). These results suggested that CFs projecting to TC2 neurons convey predictive error signals (unwarranted licks) specific to the No-go cue, in accordance with the Marr-Ito-Albus hypothesis^18, 59–61^. We postulate that cognitive error signals can be computed as sign-reversed reward prediction errors by reinforcement learning algorithms. Because TC2 activity was within 0.2 seconds, but rewards were delivered mainly between 0.5 and 2 seconds after the cue, internal forward models, which may consist of a loop network of cerebral cortex, basal ganglia and the cerebellum^68–70^, could be employed to compute the reward prediction error before actual rewards arrived^71, 72^. The proposed combination of computational reinforcement learning models and internal forward models of the cerebellum can be tested in future experiments.

Beside their correlation with error signals, synchronized CFs of TC2 neurons may also convey motor commands to control early licks in No-go trials (Fig 5C). How do climbing fibers serve a dual function of cognitive learning and motor control as reported in previous studies^53, 55^? One possibility is that PCs that are involved in cognitive learning are assumed to generate SSs that also induce licks in FA trials as motor commands. Long-term depression (LTD) may occur by co-activation of parallel fibers and CF inputs to these PCs^73–76^. As a result, simple spikes (SSs) tend to decrease and fewer erroneous licks are generated.

In contrast to TC1 and TC2 populations that exhibited marked learning effects, the TC3 population showed only slight changes in CF firing and synchrony, except for becoming silent at the final learning stage for the FA condition (Fig S5C). The correlation of TC3 CFs with reward licks was high for HIT trials across all learning stages. Indeed, TC3 neurons may be related to both motor control and sensory feedback components of reward licks since spike-triggered lick responses were large for both positive and negative time-lags (Fig S10A).

We also found that TC4 is only related to CR trials (Fig 3B and Fig S5C). Increased CF firing in TC4 neurons with learning, especially in AldC compartment 6+, might induce more successful suppression of early licks after the No-go cue, i.e., an increase of CR trials and decrease of FA trials (Fig S10B). Spike-triggered lick responses further indicated that TC4 firings suppress the licks after the No-go cue compared to those in the pre-cue period (Fig S10C). It is interesting to note that TC2 and TC4 neurons (both responded actively to the No-go cue) possess not only almost identical temporal response profiles and overlapping spatial distributions (Fig. 3), but also the largest overlap among TCs (Fig S5A). Furthermore, although fractions of TC2 and TC4 neurons changed significantly, summation of their fractions remained unchanged during the course of learning (Fig 6C). We hypothesized that TC2 and TC4 neurons are of the same neuron population, but that they changed their cue-response specificity (FA to CR) due to LTD of parallel-fiber-Purkinje-cell synapses guided by CF inputs. Let us first assume that SSs show a similar temporal profile to those of CFs (Fig 3A) for TC2 PCs with a positive baseline firing rate. At the beginning of learning, both SSs and CSs of these Purkinje cells contribute to generation of early licks, as shown by spike-triggered lick responses (Fig 5C). Once LTD occurs at parallel-fiber-Purkinje-cell synapses, SS modulation turns negative and becomes an approximate mirror image of CS modulation^77^. At later stages of learning, even though CFs of the same PCs are activated following No-go cues (Fig S5C), these neurons contribute to suppression of early licks rather than generation, because SS action is stronger than CS effects on motor control. Future simultaneous recordings of SSs and CSs as well as downstream systems may reveal neural mechanisms of these neurons.

Limitations of the current study are that activities of the four TC populations described above were of neurons sampled in different recording sessions, and that we did not investigate in detail single populations throughout learning. Thus, monitoring population activities as learning proceeds, combined with causal analysis of neuronal responses and behavior changes, is needed to understand diverse CF functions across TC populations. This is particularly crucial because CFs may multiplex motor, cognitive and reward-related information, as found in previous studies^53–56^. In Figure 6 we highlighted this possibility by showing a mixed tensor representation of 6,445 individual neurons in the cerebellar cortex. Even though each TC has a unique zonal distribution, individual zones or even neurons may represent multiple TCs and may be involved in different cerebellar functions.

From a computational learning-theory point of view, it is striking and extremely important that only four components seem to account for a wide variety of neural responses of Purkinje cells in eight AldC compartments (four TCs accounted for more than 50% of the variance of 6,445 neurons in four cue-response conditions, Fig S4A). Furthermore, these four components also explain main learning effects in behaviors, that is, more precise timing control, error decrease, reward processing, and successful lick suppression. If a learning system possesses many degrees of freedom, a huge training dataset is theoretically required. But mice learned auditory discrimination Go/No-go task within hundreds of trials. This implies that the cerebellum reduces its degrees of freedom significantly, thanks to its zonal structure and IO electrical couplings. Dimensional reduction is certainly beneficial for the cerebellum to work in concert with the cerebral cortex and basal ganglia in different learning stages^69, 78–80^.

We also found that CF synchrony is spatially and temporally dynamic, in concert with the zonal structure. This phenomenon was demonstrated by the change in the number of synchronous spikes across PCs during learning (cf. Supplemental Movies M1-2). Synchronous firing decreased in TC2 neurons for the FA, but it increased in TC1 neurons for the HIT condition. Such opposite changes in CF synchrony were strongly correlated with changes in cue responsiveness (Fig 4 and Fig S3C). This phenomenon raised an important question of which factor, CF response or synchrony, drives licking behaviors. Our decoding analysis indicated that synchronous CSs better predict licking events than ordinary CSs (Fig S8). Furthermore, previous studies showed that highly synchronous CSs caused a large drop in deep cerebellar nucleus (DCN) activity while isolated CSs produced only weak inhibition^81, 82^. These results together supported the view that synchronous CS activity shapes DCN output related to motor commands.

Remarkably, we found that CF synchrony was significantly stronger within TCs than among TCs, and the highest synchrony was observed when TCs function in their specific cue-response conditions (Fig 4A-C). Such synchrony dynamics could be realized by circuitry of Purkinje cells (PCs) - DCN – IO as follows. Under resting conditions, IO neurons are decoupled and remain rather silent due to tonic CN inhibition. When a stimulus arrives, common input activates a group of IO neurons and excites their innervated PCs, resulting in strong IO coupling due to weak CN inhibition. Cue stimulus binding with IO gap-junctions causes strong synchrony in neuronal populations that operate in the conditions in which they are engaged, e.g., TC1 in HIT trials and TC2 in FA trials. For other conditions, e.g., TC1 in FA trials and TC2 in HIT trials, the cue stimulus causes weak excitation (PSTHs of TC1-2 in Fig S5C), fails to inhibit CN inhibition, and consequently is unable to strongly couple IO neurons, resulting in weak synchrony. One may also expect that there are no common inputs across TC populations so that the electrical couplings remain decoupled among them. Together, as indicated by previous theoretical and experimental studies^15, 33, 39, 41, 46, 83, 84^, these results suggest that common inputs and positive feedback to IO cells are the key structures that organize neuron populations, induce changes in their cue responsiveness, resulting in changed behavior. Because synchronization within components is larger than synchronization across components and anatomical zones, and because even individual neurons multiplex several components, we suggest that components self-organize^85–87^ as a result of interaction between electrical synapses in the IO and positive feedback and lateral inhibition implemented by loop dynamics between PC, CN, and IO (Fig 7A).

**Figure 7:**
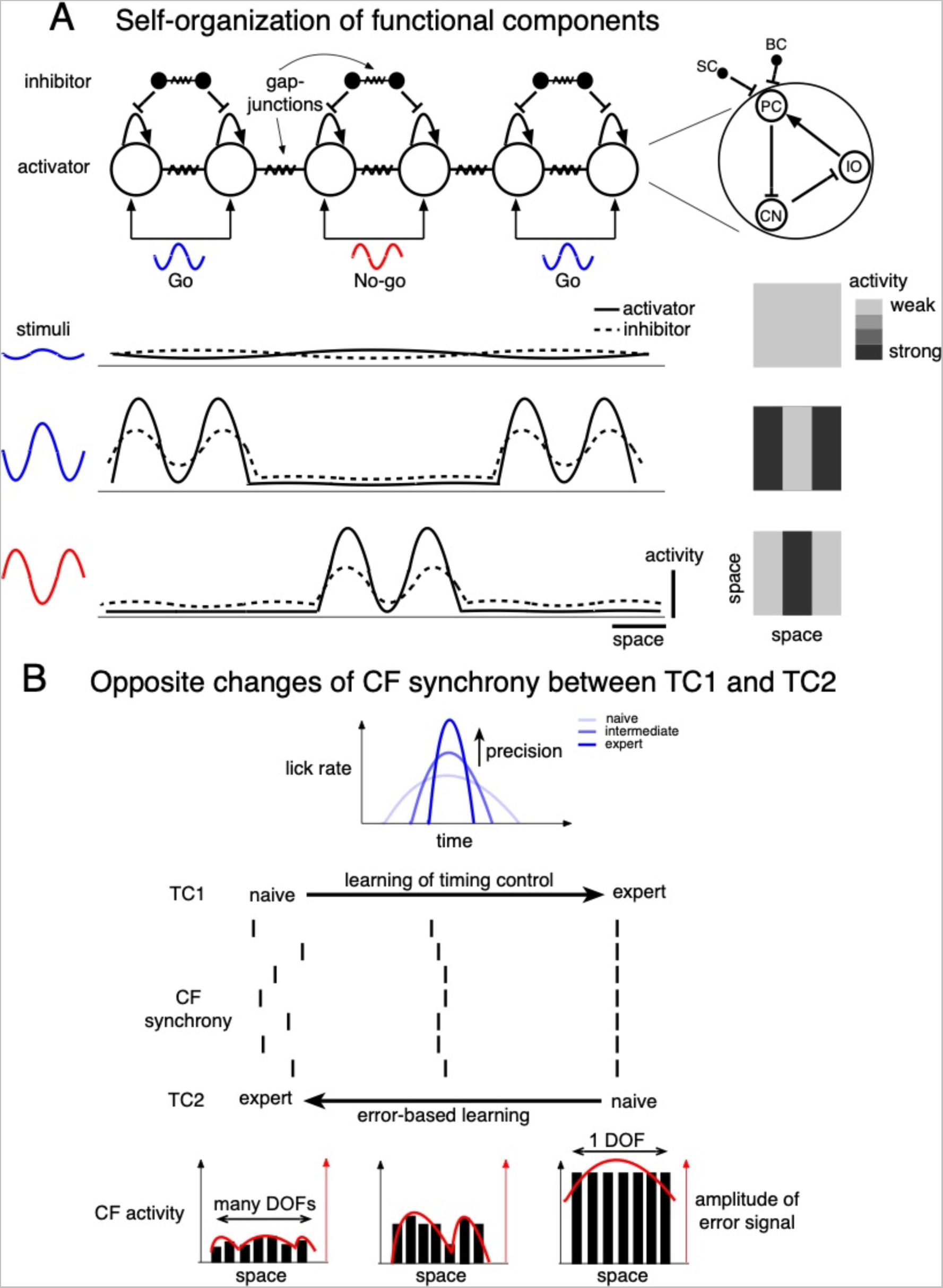
Schematic illustrations of dynamic functional organization in the cerebellum. A: Scheme for elf-organization of activator-inhibitor with diffusion system in the cerebellum. The closed-loop of IO, PC and CN could form a positive feedback circuit via double-inhibition from PC to CN and CN to IO. Also, hematic illustrations of dynamic functional organization in the cerebellum. A: Scheme for tion of activator-inhibitor with diffusion system in the cerebellum. The closed-loop ofiO. PC ld form a positive feedback circuit via double-inhibition from PC to CN and CN to 10. Also. nhibitory interneurons in the molecular layer (MLIs) inhibit themselves as well as PCs by either feedforward or lateral inhibition^89, 90^. Furthermore, both MLIs and IO neurons form densely electrically coupled neuronal networks via their gap-junctions^35, 91, 92^. Hence, cerebellar modules could self-organize their activity via a reaction-diffusion process with IO-PC-CN and MLIs acting as activator and inhibitor, respectively, and spatial diffusion is realized by their gap-junction couplings. Note that electrical interactions through gap junctions can be mathematically formulated as spatial diffusion in spatial continuity limit. When perturbed by a weak stimulus, a module maintains its stable steady state between activator and inhibitor. However, a stronger stimulus leads to promotion of the activator and destabilizes the basic state. Since the activator promotes the inhibitor, a higher activity of the inhibitor is also formed in the same region. The inhibitor inhibits surrounding activator peaks but promotes further growth of the local activator peak due to the local decrease of the inhibitor. Other activation peaks are formed by the same mechanism. The overall process could lead to formation of complicated spatiotemporal patterns, in which different spatial profiles are assumed to reflect the difference in diffusivity of the activator and inhibitor as well as stimulus strength. Here, we note that each module may selectively react to specific types of stimuli (e.g., Go or No-go cues). Thus, in the context of this study, we may regard each TC as an individual functional module, as well as a stable and non-trivial solution of the reaction-diffusion equation. B: Scheme of opposite changes in CF synchrony during learning of more precise timing control in TC1 and decreased cognitive error signals in TC2. Increased CF synchrony of TC1 caused by electrical currents flowing through gap junctions of IO neurons could stabilize timing of the synchronized motor commands and lead to more precise timing control of licks with less temporal fluctuations. By contrast, the cerebellum controls the DOF of error-based learning by synchronous CF activities of TC2 depending upon the stage of learning. In the initial stage, the error signal is large and distributed broadly in the cerebellar cortex. Strong CF synchrony with low DOF quickly reduces the error signal for fast learning. Later on, the error signal is small and distributed only within a restricted region. Desynchronized CF activity with high DOF is beneficial for more sophisticated learning.

Why does CF synchrony change in an opposite direction among neuron populations, e.g., TC1 vs. TC2? We have two possible explanations for this. One mechanistic explanation could be different synaptic plasticity rules between AldC-positive and -negative zones. Note that TC1 is mainly distributed in AldC-positive zones^9^. Another functional hint may come from the desired outputs of cerebellar coding in a particular process. Enhanced CS synchrony in TC1 would induce synchronous activation of Purkinje cell ensembles, which in turn, would tune downstream systems to facilitate initiation and coordination of precise timing control of movement. In contrast, strong CF synchrony in TC2 neurons at the initial learning stage would reduce the dimensionality of high-dimensional cognitive error signals for fast learning. At later stages, low CF synchrony would increase the dimensionality of cerebellar output for fine and sophisticated learning^15, 88^ (Fig 7B).

In conclusion, combining two-photon recordings of the mouse cerebellum in an auditory discrimination Go/No-go task with a hyperacuity algorithm of spike-timing estimation and tensor-component analysis, we demonstrated that motor and cognitive functions were learned in distinct neuronal populations. Compartmental representations of these populations align well with functional and anatomical boundaries between medial and lateral parts of the cerebellar hemisphere, as well as expression of AldC. Furthermore, CF synchrony is a plausible mechanism to induce changes in both cue responsiveness and compartmental representation along with learning. Bidirectional synchronous response-associated changes in CF activities finely constructed on compartmental structure could provide a flexible learning scheme in diverse cerebellar functions.

## Methods

All experiments were approved by the Animal Experiment Committees of the University of Tokyo (#P08-015) and the University of Yamanashi (#A27-1). They were conducted in the same way as reported in Tsutsumi et al. (2019)^50^. Briefly, 17 adult male heterozygous Aldoc-tdTomato mice^49^ and adult male wild-type mice (Japan SLC, Inc, n = 5) at postnatal days 40–90 were used. A cranial window was created over the left Crus II of mice anesthetized with isoflurane (5% for induction). Two-photon imaging was performed during the following Go/No-go task phase at a scanning rate of 7.8 Hz, using a two-photon microscope (MOM; Sutter Instruments) equipped with a 40x objective lens (Olympus) controlled by ScanImage software (Vidrio Technologies. Imaging data were analyzed using MATLAB (R2018a; MathWorks). Two-photon recording experiments were conducted once daily for 4-7 consecutive days, for a total of 236 sessions, each containing 13-43 trials.

### Two-photon recordings of climbing fiber responses

Each trial with two-photon recording data was categorized as HIT, FA, CR, or MISS, according to licking behavior within a response period of 1 s to the two cues, i.e., correct lick in response to the go cue, unwarranted lick in response to the No-go cue, correct response rejection to the No-go cue, or response failure to the Go cue, respectively. Each session was also categorized according to the fraction correct of mouse performance, i.e., the ratio of correct responses (HIT and CR) to all trials of the session. Categories included the 1st, 2nd, and 3rd stages of learning with the fraction correct <0.6, 0.6 - 0.8, and >0.8, respectively. The fraction correct for the Go cue was evaluated as the ratio of HIT trials to Go trials. The fraction incorrect for No-go cue was evaluated as the ratio of FA trials to No-go trials (Fig 1C). For each session, on average, there were approximately 30 neurons simultaneously recorded while more than 30 trials of Go/No-go cues were randomly presented. In total, there were 59, 83, and 94 recording sessions, 1462, 2405, and 2578 neurons and 1552, 2731, and 3692 trials in the 1st, 2nd, and 3rd stages, respectively (see Tables S1-2 for more details).

### Calculation of licking variables associated with cues

We evaluated two licking variables associated with learning of Go and No-go cues in the early response window (0 – 0.5 s after cue onset). Lick-latency was estimated as the difference in timing of the first lick onset and the cue onset. We found that, during learning, there were no systematic changes in the mean of lick-latency, but its fluctuation from the mean was significantly reduced (Fig S1A). To account for slight variations in lick-latency across training sessions, we fitted the lick-latency as a function of the trials, sorted by training session, by a 4^th^ order polynomial curve for individual mice, which is the best model among 0 to 5th order polynomial models with Akaike Information Criterion. The lick-latency fluctuation of a single trial was then calculated as the absolute difference between the lick-latency and the fitted curve, normalized by the mean of lick-latency for individual mice (Fig S1B). Lick-latency fluctuation was computed only for HIT and FA trials, omitting MISS and CR trials, for which there were no licks in the early response window. The early lick rate was counted as the number of licks in the early response window. For each animal, we fitted lick-latency fluctuation in Go and number of early licks in No-go trials using two linear models of trials, respectively; hence, their slopes indicated the corresponding learning effects, i.e., negative slopes for less fluctuation in lick-latency after Go cue and less erroneous licks after No-go cue.

### Reconstruction of spike events

Ca signals in Purkinje cell dendrites were evaluated for regions-of-interest (ROIs) of the two-photon images selected by Suite2p software^93^. Spike trains were reconstructed for 6,445 Purkinje cells sampled in the 17 mice, using hyperacuity software^51^ (HA_time) that detected spike activities for Ca signals of two-photon imaging with a temporal resolution of 100 Hz. The mean CF firing rate (1.1 ± 0.4 spikes/s) and cross-correlograms (CCGs) of CFs across neurons within individual Ald-C compartments were consistent with those for previous studies using electrical recordings in behaving^65^ and anesthetized^35^ mice (Fig S2B&C). Furthermore, simulations of observed Ca signals using GCaMP6f dye showed that the HA_time was capable of detecting roughly 90% of the spikes (Fig S2D). Together, these results guaranteed the reliability of HA_time in detecting CF spike timing with high temporal precision. We note that to compensate for small jitters of spike timing estimation as well as to increase the signal-to-noise ratio, we used 30-ms bins for evaluating the synchrony and 50-ms bins for constructing the peri-stimulus time histograms of CF activity (see below).

### CF responsiveness to cue stimulus

For evaluating the CF responsiveness of AldC compartments to the cue stimulus, we constructed the peri-stimulus time histogram (PSTH) of CF activity with a time bin of 50 ms. PSTHs were constructed for individual Purkinje cells and averaged across Purkinje cells in the same AldC compartment (Fig 2A-D). We also evaluated response strength as cumulative PSTHs for 0-300 ms after cue onset, since the peaks of PSTHs were mostly 200-300 ms. To demonstrate opposite response changes between HIT and FA of medial and lateral parts of Crus II, for each neuron, we computed the difference in response strengths between HIT and FA trials (i.e. HIT - FA). We then selected the top 100 neurons in each AldC 5+ and 6- with highest and lowest values, respectively.

### Synchrony analysis of CF activity

In this study, we evaluated synchronization of CF activities by two indices, one on a trial basis (we named it “instantaneous synchrony”) and the other across trials in the same recording session (“synchrony strength”). For the former, we estimated the instantaneous synchrony in each trial by the total number of synchronous spike pairs (co-activated in time bins of 30 ms) in the window of 300 ms before the first lick, normalized by the number of cell pairs. Suppose that 10 neurons were simultaneously measured and together they produced 20 spikes within the time window. Further suppose that 4 neurons are co-activated in one time bin and 2 neurons were co-activated in another time bin (the other 14 spikes fired in separated time bins). Then instantaneous synchrony was calculated as 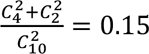. For this example, the spike count was 20/10 = 2. We note that for the CR and MISS trials, for which no early lick was generated, the time window was fixed as 0-300 ms after cue.

We also measured the synchrony strength across trials in two different neurons by the cross-correlogram. The spike train of a neuron was represented by *X(i)*, where *i* represents the time step (*i = 1, 2, …, N*). *X(i) = 1* if spike onset occurs in the *i-*th time bin; otherwise, *X(i) = 0*. *Y(i)* was the same as *X(i)*, but for the reference neuron. The cross-correlation coefficient at time lag *t*, *C(t)*, was calculated as follows:

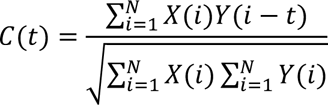

A 10-ms time bin was used; thus, for two spikes to be considered synchronous, their onsets must occur in the same 10-ms bin. Synchrony strength was defined as the sum of *C(t)* in a window of ±10 ms around the zero-lag time bin *C(0)*. Cross-correlation caused by spike synchronization to the task event stimulus was evaluated as a CCG, for which the spike time of the reference neuron was shifted by one trial period^94, 95^.

### Tensor component analysis

Since most CFs reduce their activity to baseline 2 s after the auditory cue, we conducted tensor component analysis^52^ on PSTHs sampled from −500 ms to 2 s from cue onset (bin size, 50 ms) of all Purkinje cells (*n* = 6,445) in the four cue-response conditions, i.e., HIT, FA, CR, and MISS. Each PSTH was subtracted from its baseline activity, defined as the mean value of the PSTH in the range of [−2, −1] s before cue onset. We note that if the number of trials of a specific cue-response condition in a recording session is less than 5, we fix the corresponding PSTHs of the neurons in that session to 0.

Let x_ntk_ denote the PSTH of neuron *n* at time step *t* within cue-response condition *k*. TCA yields the decomposition

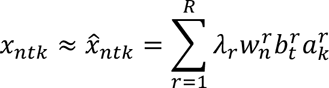

where *R* is the number of tensor components, 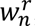, 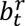 and 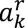 are the coefficients of the neuron, temporal, and response condition factors, respectively. Those coefficients were scaled to be unit length with the rescaling value 𝜆*_r_* for each component *r*. We introduced a non-negative constraint of those coefficients (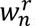 ≥ 0, 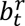 ≥ 0 and 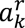 ≥ 0 for all *r*, *n*, *t* and *k*). Figure 3 showed the coefficients [inline1]and [inline1] for each of the tensor component *r*=1,2,3,4.

TCA iteratively estimated the coefficients with an alternating least-squares algorithm; thus, its results are dependent on random initial values. The number of tensor components was systematically examined with *R*=1-10, each with 100 random initializations. For each *R*, we selected the optimal solution as the one that was obtained most frequently among 100 initializations. To evaluate the fitting performance, the original PSTHs, *x*, and those reconstructed from TCA coefficients, 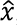, 0-1 second after cue onset were first low-pass filtered (cut-off frequency of 2 Hz). Then variance accounted for (VAF) was computed as follows

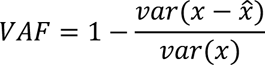

For each value of *R*, we further inspected the similarity of the optimal solution and the solutions obtained from the other 10 random initializations. We selected *R*=4 which accounted for more than 50% of total variance and provided stable solutions (see Fig S4A-B). We noted that increasing *R*=1-3 to 4 added meaningful components, i.e., segregation of TC4 from TC2 specific for CR trials. However, further increasing *R* separated the components with slight differences observed only in temporal profiles (Fig S4C).

### Sampling neurons by TCA coefficients

For further detailed analyses of the four tensor components retrieved by TCA, we sampled neurons that were best represented by each component as follows. First, at each learning stage (1st - 3rd stages), we selected the top 300 neurons that had the largest coefficients for each of the four components. Then, neurons that were sampled by more than one component, i.e., that overlapped, were excluded (Fig S5A). As a result, we sampled 2,096 neurons for the four tensor components. Note that, because different numbers of neurons were recorded across learning stages (Table S1), we also sampled the top 10-20% of TC neurons using the above process and found identical PSTHs of TCs (Fig S5C).

### Spike-triggered lick response

To investigate the correlation of CF activity and licking behavior, we sampled spikes and lick onsets in the three windows of each trial: 0-0.5 second (early lick), 0.5-2 second (reward lick) and 2-4 second (succeeding lick) after cue onset. Then, the spike-triggered lick response was constructed across trials with a time bin of 100 ms, normalized by the total number of trials. We note that for TC1 and TC2 (c.f. Fig 5), we examined lick responses for synchronous spikes, those that were co-activated (in the time bin of 30 ms) with at least one spike of the other neurons of the same TC in the same recording session. For TC3 and TC4, since their synchrony changes were modest (Fig S6), we examined all sampled spikes (Fig S10).

### Regression analysis of synchrony and licking variables

We fitted a multiple linear regression model on a trial basis, with the synchrony of each TC as an explanatory variable and two licking variables as response variables. The formula of this model specification was shown by Wilkinson notation as

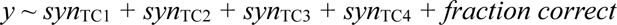

in which *y* was either lick-latency fluctuation or the number of early licks, *syn*_TC1-TC4_ were synchrony of TC1-4 neurons of the same trial, and *fraction correct* was common across trials in the same recording session. *fraction correct* was introduced to represent general learning effects, except neural synchrony. Note that the variable *syn*_TC1-TC4_ of a particular trial was set to 0 if there were no selected neurons as TC1-4 in that trial. For example, suppose that there were 2 and 4 neurons selected as TC1 and TC2 for a single trial, respectively. Then *syn*_TC1_ and *syn*_TC2_ were computed as the number of synchronous spikes of 2 and 4 neurons for that trial, respectively, and *syn*_TC3_ = *syn*_TC4_ = 0. Due to the neural sampling process, there existed 2,465 trials for which none of the neurons was selected by any of the four TCs. We excluded those trials from the analyses. As a result, the multiple regression was conducted for 3,080 trials. We note that the linear mixed-effects model with random effects for intercept grouped by animal, i.e, *y ∼ syn*_TC1_ *+ syn*_TC2_ *+ syn*_TC3_ *+ syn*_TC4_ *+ fraction correct + (1|animal)*, with *animal*=1..17 as the mouse index, showed little difference from the above model, indicating that there was no across-mice effect.

We constructed added variable plots, in which variables were adjusted for visualizing partial correlations between licking response variables and an individual explanatory variable (predictor) conditional on other explanatory variables (c.f. Fig 5 and Fig S7). Adjusted values are equal to the average of the explanatory variable plus the residuals of the response variable reconstructed by all explanatory variables except the selected explanatory variable. Note that the coefficient estimate of the selected predictor for the adjusted values is the same as in the full model that includes all predictors. Multiple regression fitting and added variable plots were conducted using MATLAB functions, *fitlm* and *plotAdded*, respectively.

### Statistics

All statistical analyses were performed using MATLAB software. Unless otherwise stated, data are presented as means ± SD. The non-parametric Kruskal-Wallis one-way analysis of variance test was used to determine whether data groups of different sizes originate from the same distribution. Significance level: n.s, p > 0.05; * p < 0.05; ** p < 0.01; *** p < 0.001; **** p < 0.0001.

## Acknowledgments

H.H., K.K. and K.T. were supported by Grants-in-Aid for Scientific Research in Innovative Areas (17H06313) and for Transformative Research Areas (22H05160 to M.M., 22H05156 to M.Kawato, and 22H05161 to K.K.). H.H. and K.T. were partially supported by JST ERATO (JPMJER1801, “Brain-AI hybrid”). H.H., M.Kawato, and K.K. were partially supported by the Grant Number JP21dm0307002, JP21dm0307008, and JP19dm0207080, respectively, Japan Agency for Medical Research and Development (AMED). M.Kawato was partially supported by Innovative Science and Technology Initiative for Security Grant Number JP004596, Acquisition, Technology & Logistics Agency (ATLA), Japan. M.Kano and K.K. was partially supported by Grants-in-Aid for Scientific Research (JP18H04012, JP20H05915, JP21H04785 to M.Kano and JP22H00460 to K.K.) from Japan Society for the Promotion of Science (JSPS).

## Supplemental Information

### Examination of the hyperacuity algorithm (HA_time) by simulations

We examined HA_time by simulating Ca signals that reproduce those observed in two-photon recordings. In particular, spike events were generated according to a Poisson distribution with a mean firing rate of 1 Hz. Ca responses were simulated by convolving the double exponentials with the spike events. The rise time constant was fixed at 10 ms, while the decay time constant was 300 ms corresponding to that of the GCaMP6f dye. Gaussian noise was added to reproduce the SNR (SNR = 10 dB) of experimental data. A total of 5 cells each with 100 spikes were generated for testing HA_time performance (Fig S2).

For performance evaluation, a correct hit case was defined as one in which the time difference between an estimated spike and a true one was smaller than the sampling interval (100 ms in simulations). This process was repeated to find all hit cases between two spike trains. Remaining spikes in the true spike train were counted as missed spikes while spikes remaining in the estimated spike train were false-positives. We used the f1-score, the arithmetic mean of the sensitivity and precision, to evaluate spike detection performance.

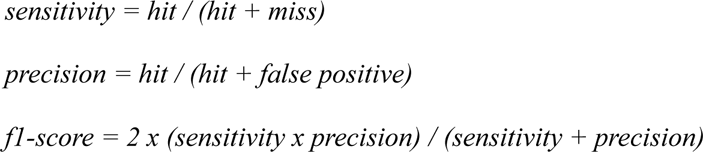

### Decoding analysis

We conducted a decoding analysis to test whether synchronous spikes of TC1-2 neurons could predict the occurrence of lick events in the 0-1s window better than the other three models - including all spikes of TC1-2 neurons, all spikes from all neurons in the same recording session, and a “chance” model for which there was no spike-lick correlation.

We first constructed a common spike-triggered lick response for the entire spike and lick events in HIT and FA trials (Fig S9A). From a probabilistic view-point, this spike-triggered lick response could be considered as the probability of a lick event *l* given a single spike *s* in the same trial, *p(l | s)*. With an assumption that a single spike caused a lick independently, the probability of a lick event given a spike collection *S^M^* of the model M follows:

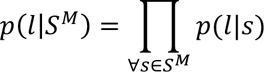

for the chance model independent of spike sequences, *p(l | S^M^) = 1/dt_0_*, where *dt_0_* is the sampling rate of licking events (Fig S9B). Finally, the total log-likelihood of the model *M* was computed for the entire licking events *L* as

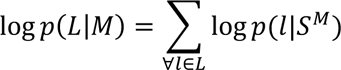

As a result, synchronous spikes of TC1-2 predicted occurrence of lick events (total log-likelihood for all lick events, −32541 and −7603 for TC1 in HIT trials and TC2 in FA trials, respectively, Fig S8) statistically better than all spikes of TC1-2 neurons (−32825 and −7657), all spikes of all neurons in the same recording session (−33494 and −7751) and the chance level for which no correlation between spike and lick events was assumed (−35312 and −8146). Note that the probabilities were trial-wise and that all lick and spike events were sampled in 0–1 second after cue onset.

## Supplemental Figures

**Figure S1:**
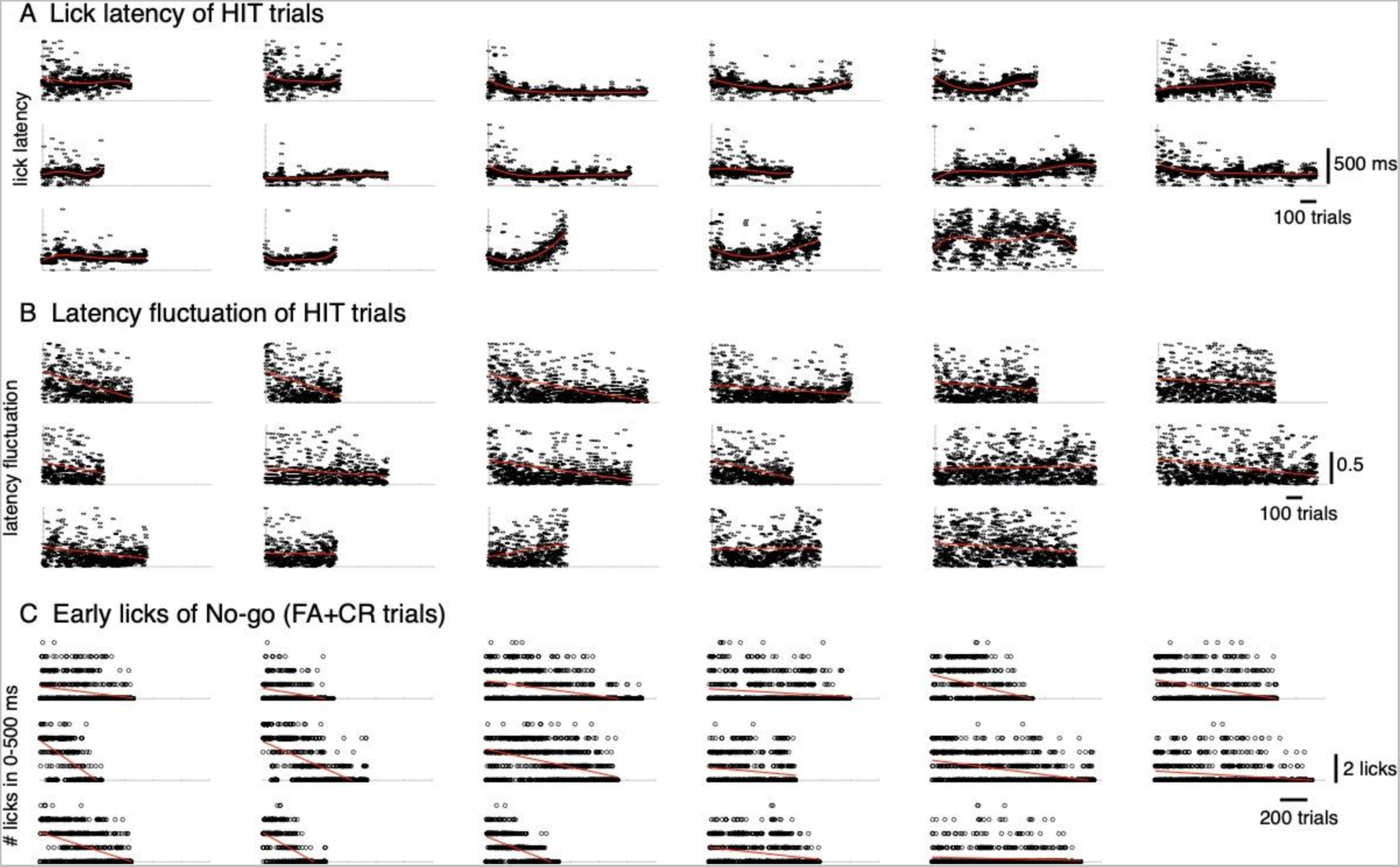
Licking behavior in the early response window for individual mice. The lick-latency (A) and lick-latency fluctuation (B) in HIT trials, early licks in No-go (CR and FA) trials, estimated from a window of 0-500 ms after cue, of 17 mice. Trials were sorted by training session. The red traces in A-C indicated polynomial fits of the variables as the functions of trials (4^th^ order for A and 1^st^ order for B and C).

**Figure S2:**
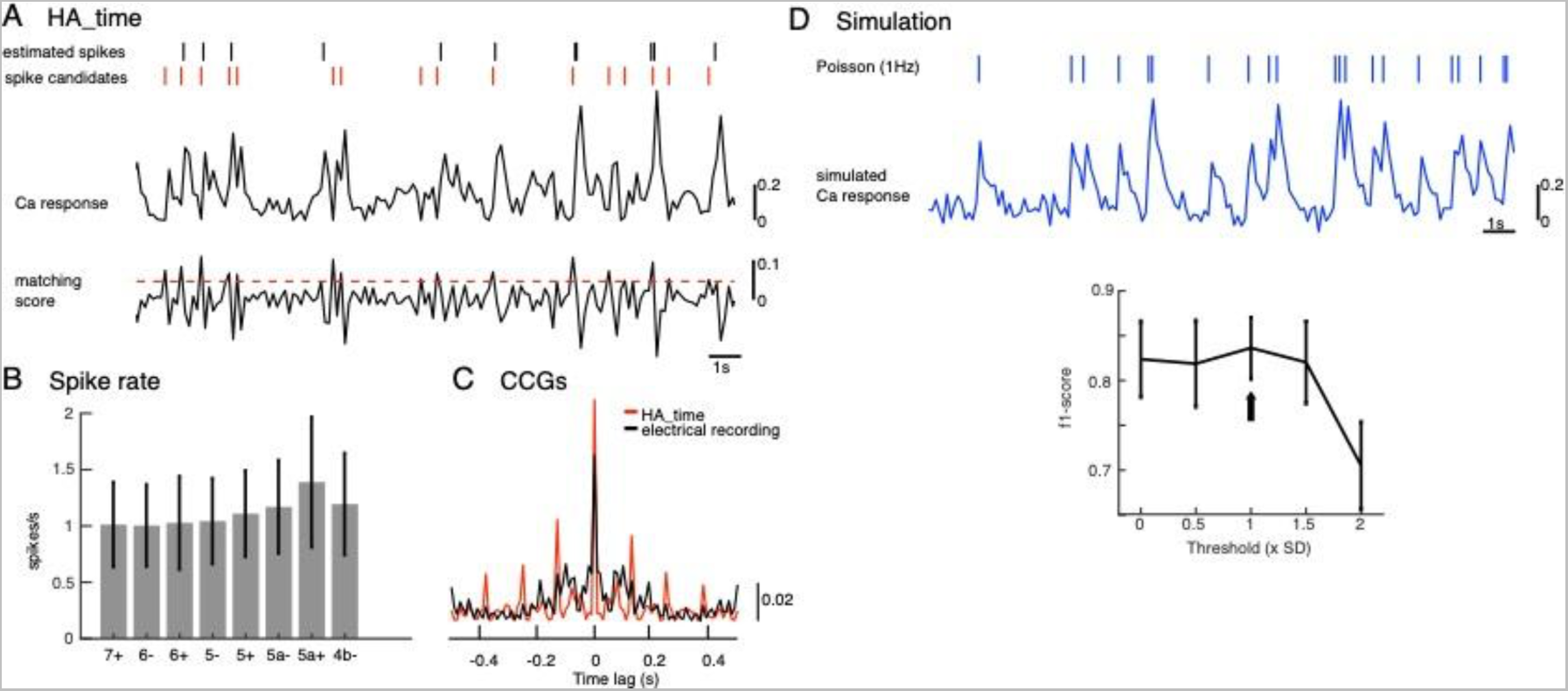
Illustration of CF reconstruction by HA_time and its examination. A: The spike model (inset) was estimated for Ca signals (top trace) of two-photon recordings by Bayesian inference, assuming size and shape constancy of spikes. Spike candidates (red short bars) were selected by thresholding (threshold = 1 SD, red dashed line) matching scores of Ca signals (bottom trace) with the spike model, and spikes were selected by SVM from spike candidates. We optimized the threshold for matching score so as to maximize the f1-score so that it was >0.8 (black arrow Fig. S2D). Finally, spike timings (black short bars) were estimated with temporal resolution of 100 Hz, so as to minimize residuals of observed and predicted Ca signals by the model^51^. B: spike rates of estimated CFs by HA_time across the AldC compartments. The mean firing rate (1.1 ± 0.4 spikes/s) agrees with electrical recordings in behaving mice^65^. C: population CCG of spikes estimated by HA_time in a recording session of the AldC compartment 5+ (red histogram, n = 20 cells) is consistent with that of multichannel electrical recording of Purkinje cells (black histogram, n = 25 cells) reported in Blenkinsop and Lang (2006)^35^. D: Upper: simulation of the Ca response (blue trace) was generated by convolution of the spike model and Poisson spikes (rate, 1 Hz, short blue bars), adding Gaussian noise (SNR = 10). Lower: f1-score estimated by HA_time with the threshold values varied from 0 to 2SD in spike candidate selection. Simulated Ca responses of a total of 5 Purkinje cells indicated that HA_time was capable of detecting roughly 90% of the spikes.

**Figure S3:**
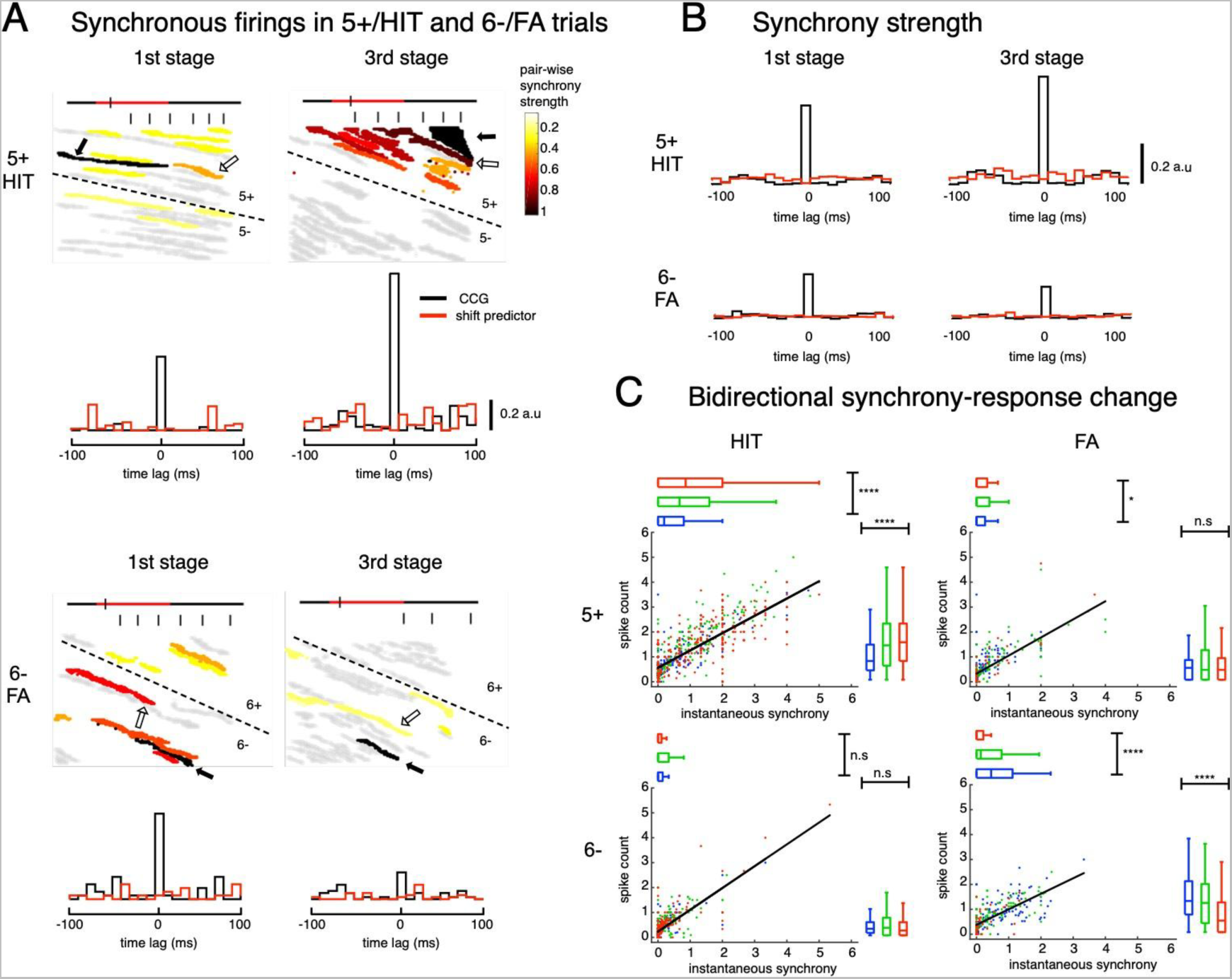
Synchrony dynamics and associated synchrony-response bidirectional changes in AldC compartments 5+ and 6-. A: synchrony analysis of representative sessions for 5+/HIT and 6-/FA trials. Upper: snapshots of synchronous firings in HIT trials at the boundary 5+/5- of two representative sessions in 1st and 3rd stages. Reference cells (black arrows) are those that show the largest difference in HIT-FA responses. Stair-plots indicate CCGs between the reference cell and the proximal cell (open arrows) that has the highest pairwise synchrony strength in the session. Lower: similar plots of representative sessions for 6-/FA trials. B: population CCGs for all sampled cell pairs in 5+/HIT (upper) and 6-/FA trials (lower) for the 1st and 3rd stages. Red traces in CCGs indicate shift predictors estimated for the correlation solely to cue stimuli. C: scatter plots of averaged spike counts and instantaneous synchrony across all trials, both estimated in the window of 300 ms before the first lick, for HIT (left column) and FA (right column) trials in three stages of learning for two representative AldC compartments 5+ (upper row) and 6- (lower row). Thick black trace indicates correlation of the two quantities. Summary statistics of instantaneous synchrony and spike count were shown by horizontal and vertical box plots, respectively. For each box plot, the bar indicates the 25% and 75% and the central mark indicates the median. The whiskers extend to the most extreme data points not considered outliers. Asterisks indicate significant level of one-way ANOVA: n.s, p > 0.05; * p < 0.05; **** p < 0.0001.

**Figure S4:**
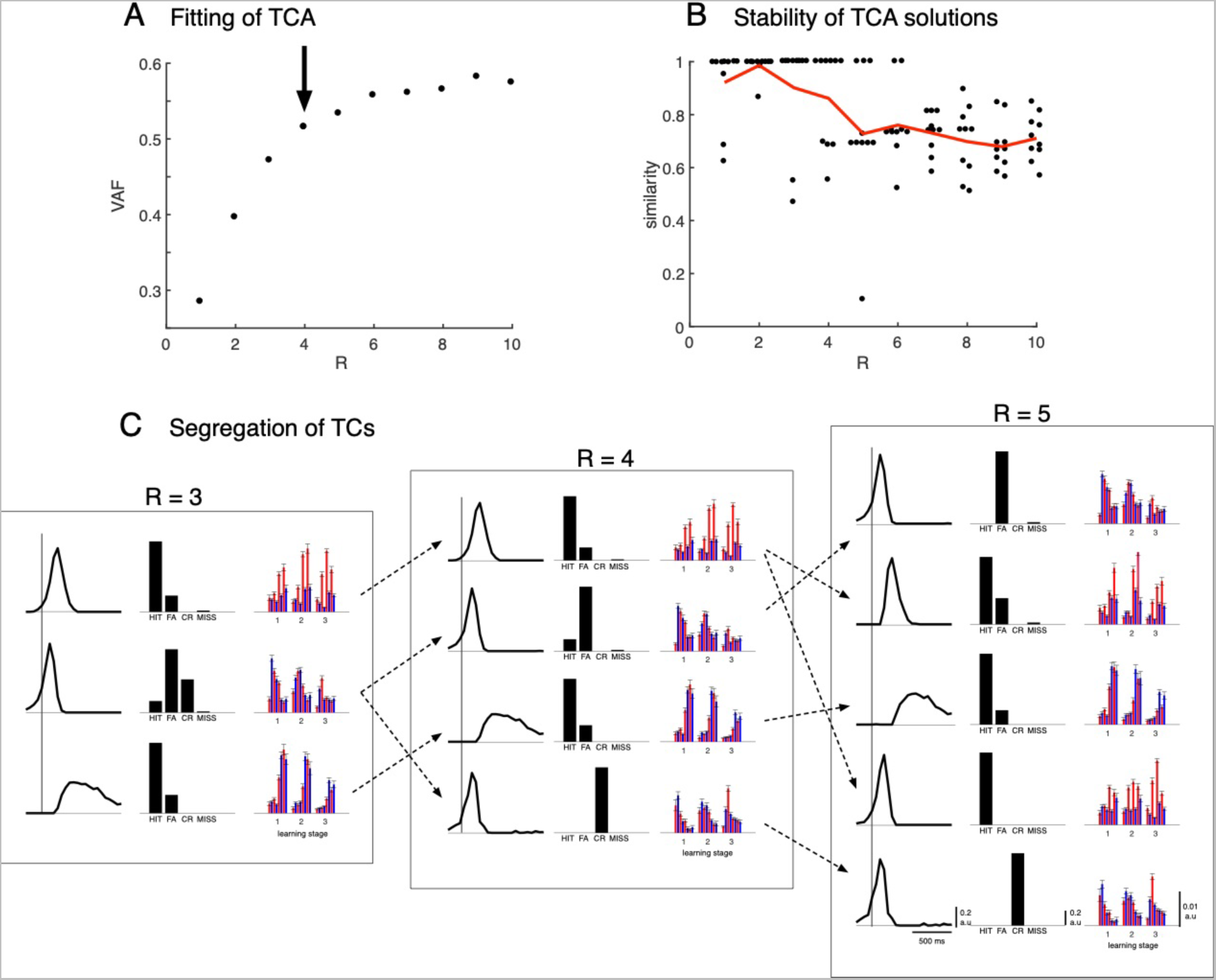
Tensor component analysis of population PSTHs with a varied number of tensor components. A: the fitting performance of TCA, which estimates variance accounted for (VAF) of the reconstructed data using the optimal solutions (see Methods for details), with the number of tensor components varied from R=1-10. Arrows indicate values at R = 4. B: similarity scores, which measure the similarity between solutions of 10 random initializations (black dots) with the optimal solution, for each of the number of tensor components R. C: optimal TCA solutions for R = 3, 4, 5 showed the segregation of TCs while increasing R. Dashed arrows were drawn by visual inspection of the similarity in temporal profile, cue-response condition and zonal distribution, between the TCs.

**Figure S5:**
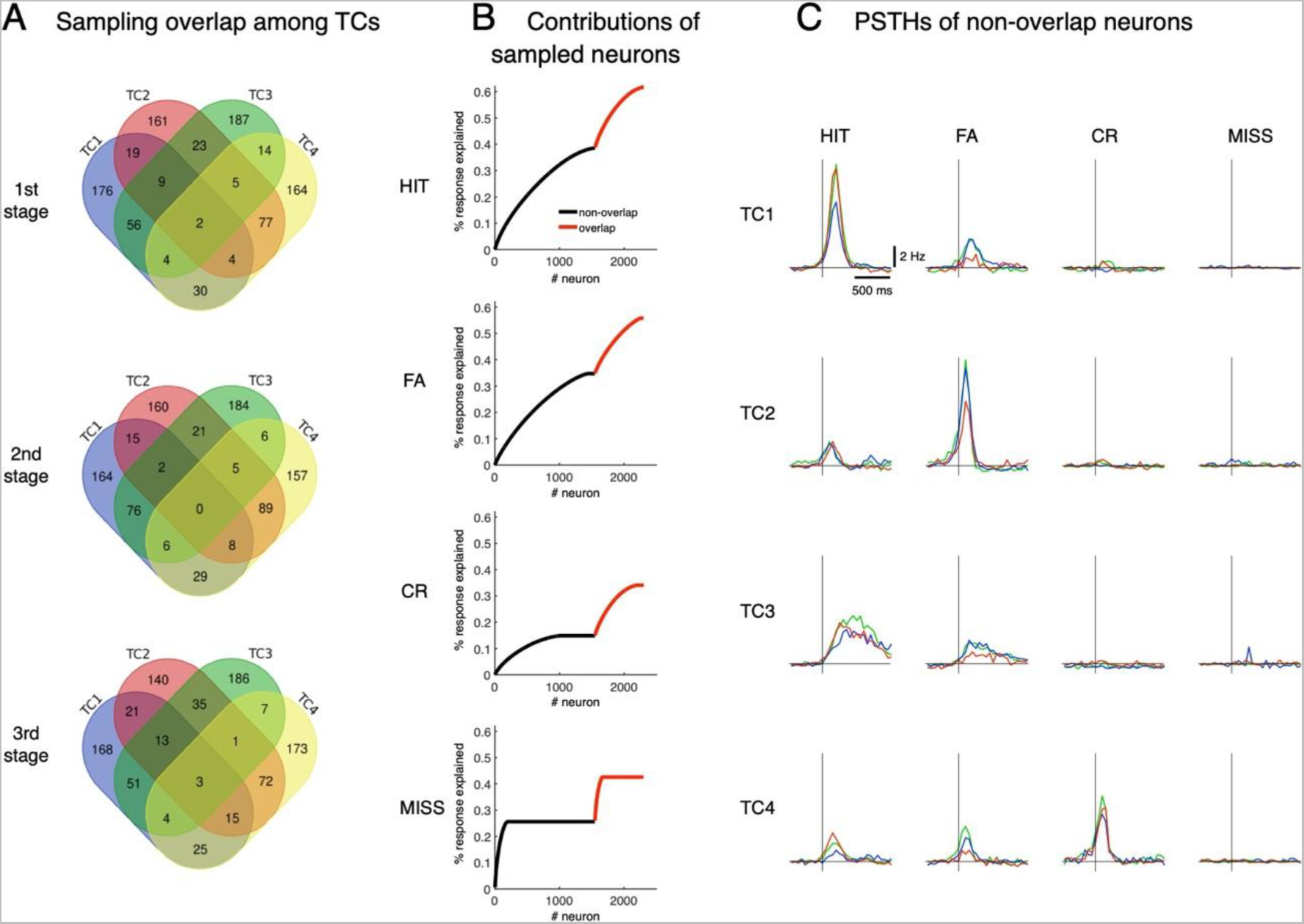
Neural sampling by TCA. A: At each learning stage, we sampled 300 neurons whose coefficients were highest for each of the tensor components, TC1-4. The Venn diagram shows the number of overlapping neurons by this sampling. We excluded overlapping neurons from further analysis in the main results. B: The response explained was estimated by the ratio of accumulated responses of sampled neurons for 1 s after cue onset to the accumulated response of a total of 6,445 neurons. Black and red are for selected neurons without overlapping ones and overlapping neurons. C: PSTHs in the four cue-response conditions of selected neurons without overlapping neurons.

**Figure S6:**
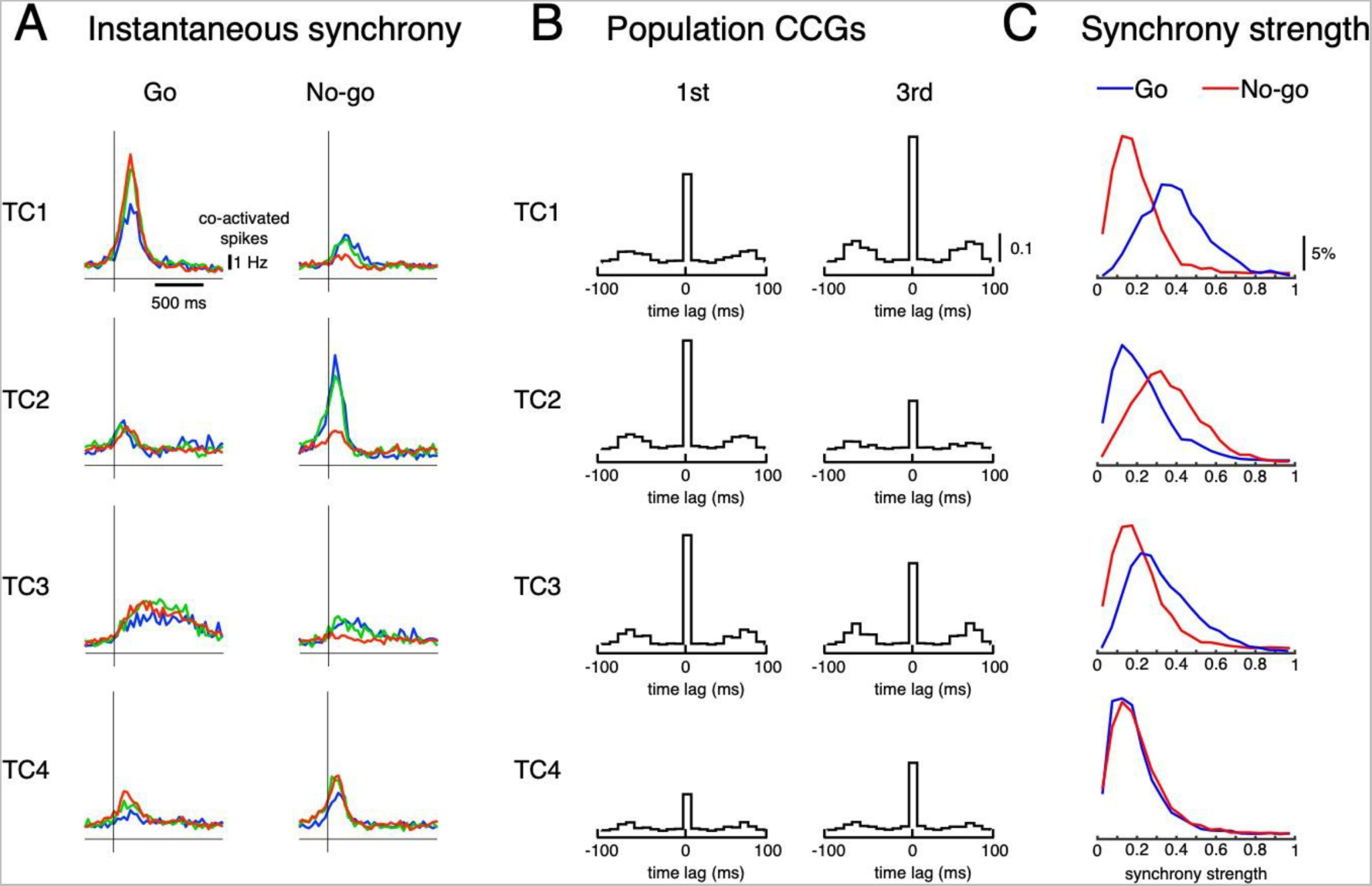
Synchronous firings of TCs. A: averaged number of co-activated neurons following Go and No-go cues of the four TCs. Averaged instantaneous synchrony across trials. B: population CCGs of sampled TC neurons in the 1st (first column) and 3rd (second column) learning stages. C: histograms of synchrony strength of sampled TC neurons for Go (blue) and No-go (red) cues. These results indicate that synchrony was high in cue-response conditions that are maximally associated with TC activities.

**Figure S7:**
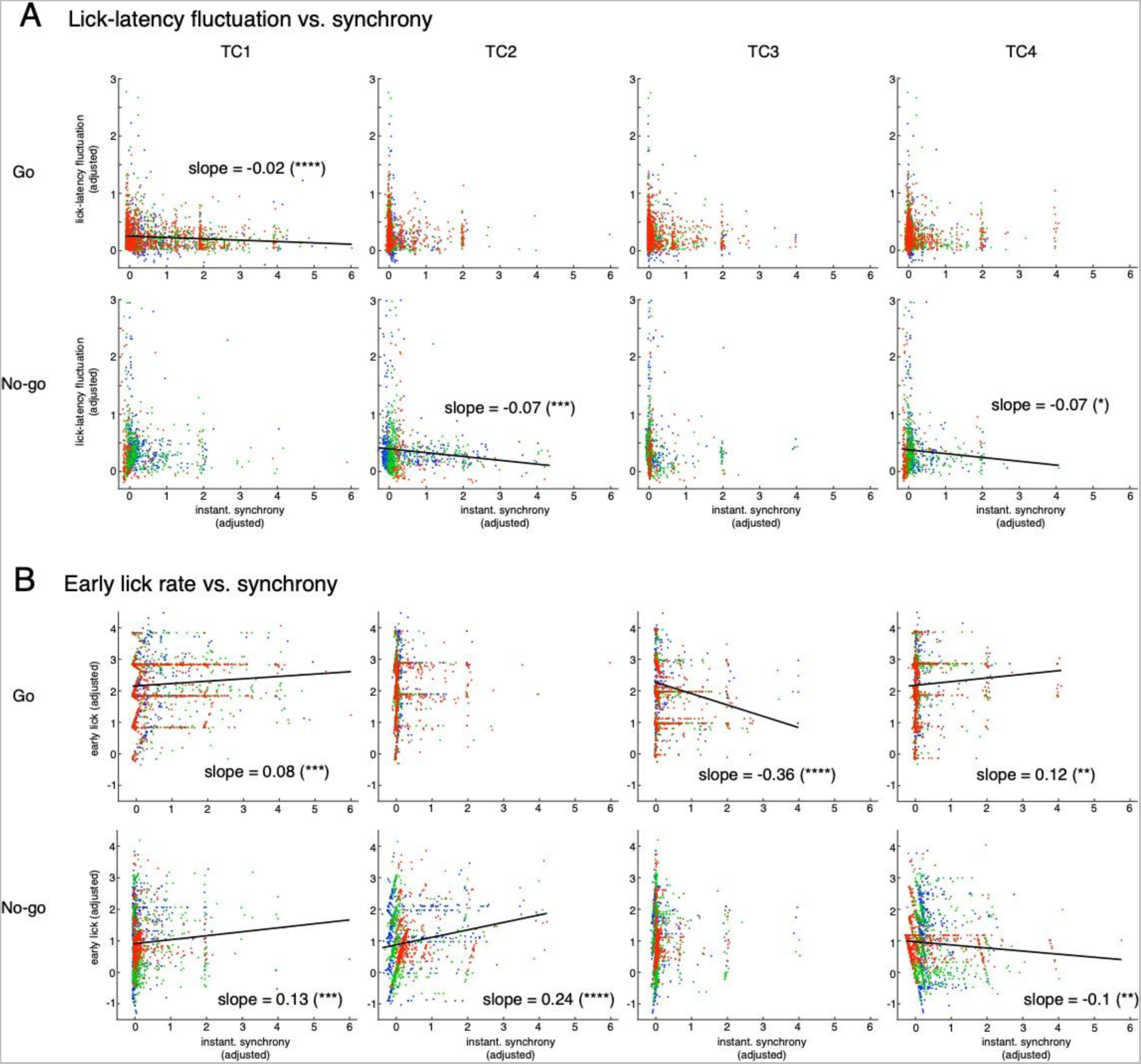
Multiple regression analysis for two lick variables. (lick-latency fluctuation – A and early lick rate – B) and synchrony of TC1-4 (see Methods for details). The top and bottom rows in A and B are for Go and No-go cues, respectively. The four columns are for TC1 – TC4 neurons. Only significant correlations (p < 0.05) are shown by black lines with slope values. The ordinate and abscissa were adjusted to show partial correlations of lick variable and synchrony of single TCs (see Methods for more details). Note that the slopes for fraction correct variable were all significant (p < 0.0001), but negative for lick-latency fluctuation/Go (top row in A) and early-lick-rate/No-go (bottom row in B) and positive for lick-latency fluctuation/No-go (bottom row in A) and early-lick-rate/Go (top row in B) combinations. These results are consistent with the behavioral results that there was no learning in reduction of lick-latency fluctuation for No-go trials or early lick rate for Go trials.

**Figure S8:**
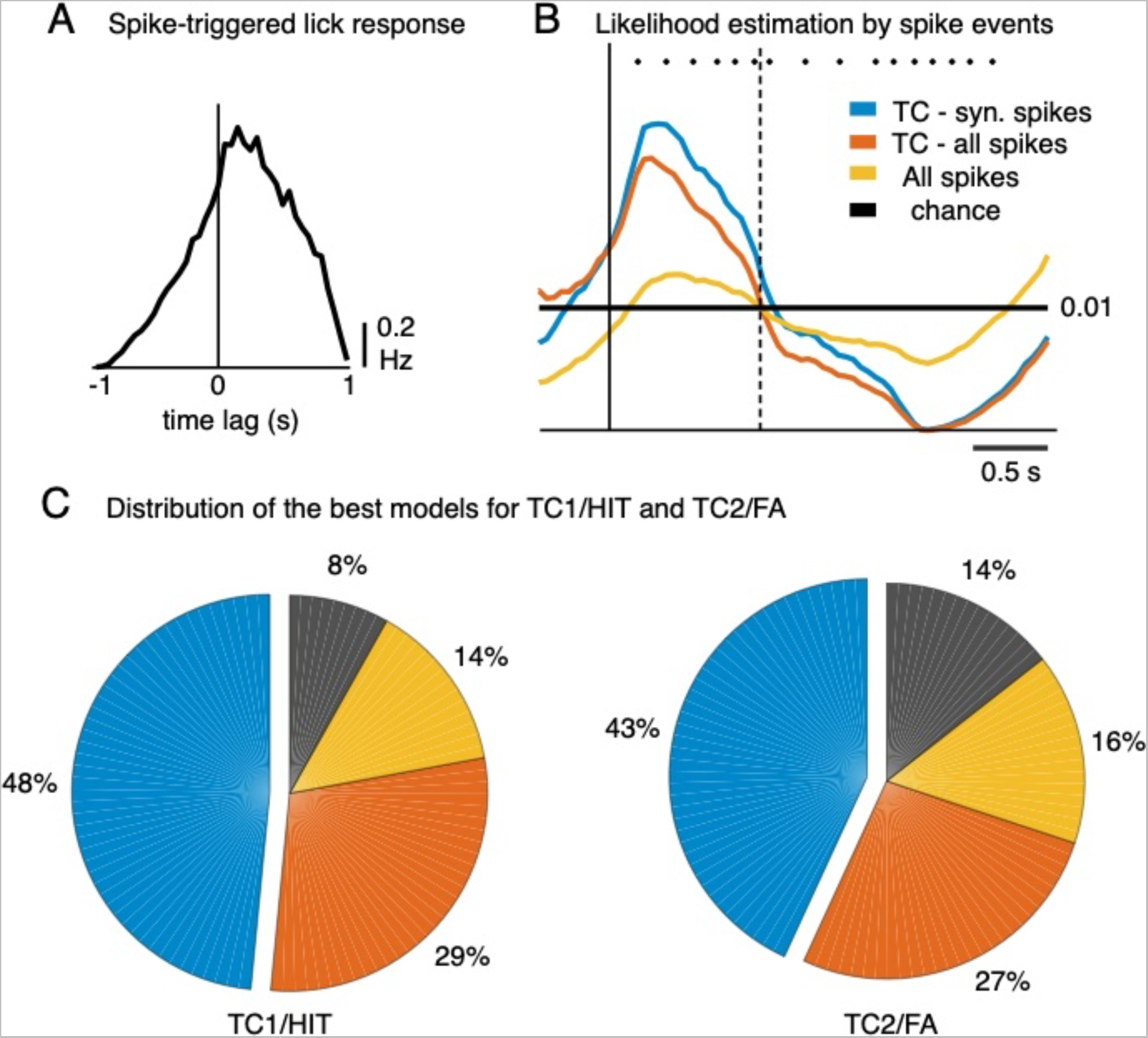
Decoding analysis of lick events. A: spike-triggered lick response for entire spike and lick events sampled from 0-1 second after the cue onset in HIT and FA trials. B: Likelihood estimation of a lick given different kinds of spike events in a representative HIT trial. Spike events were sampled according to the three spiking models, including synchronous spikes of TC1 neurons (blue), all spikes of TC1 neurons (orange), and all spikes of all neurons in the same recording session (yellow). Note that 0.01 indicates the chance level (black) to correctly predict the occurrence of a single lick at 100-Hz precision. Black dots indicate licking events. C: For each single lick event, the best model was determined for the maximal likelihood among the four models. Pie charts indicated the percentage of the best model for the entire TC1/HIT (left) and TC2/FA (right) trials.

**Figure S9:**
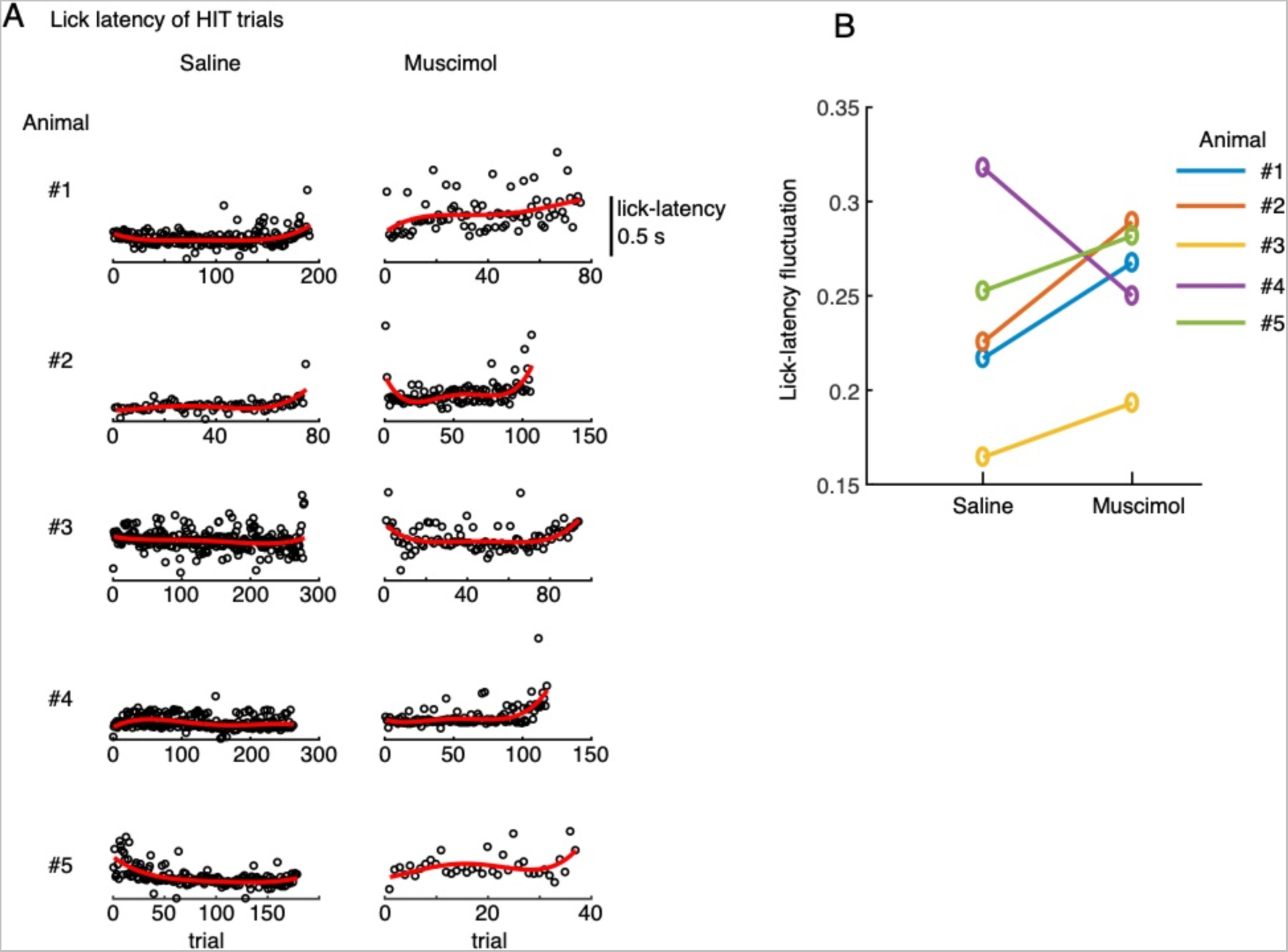
Effect of muscimol injection on lick timing precision. A: Lick-latency in HIT trials for saline (left) and muscimol (right) conditions of 5 animals. Red traces showed 4^th^ order polynomial fits of the lick-latency as functions of trials. B: increases of mean lick-latency fluctuation (ordinate, see Methods for more details) in 4 out of 5 animals indicated that muscimol effectively reduced the precision of lick timing compared with saline conditions.

**Figure S10:**
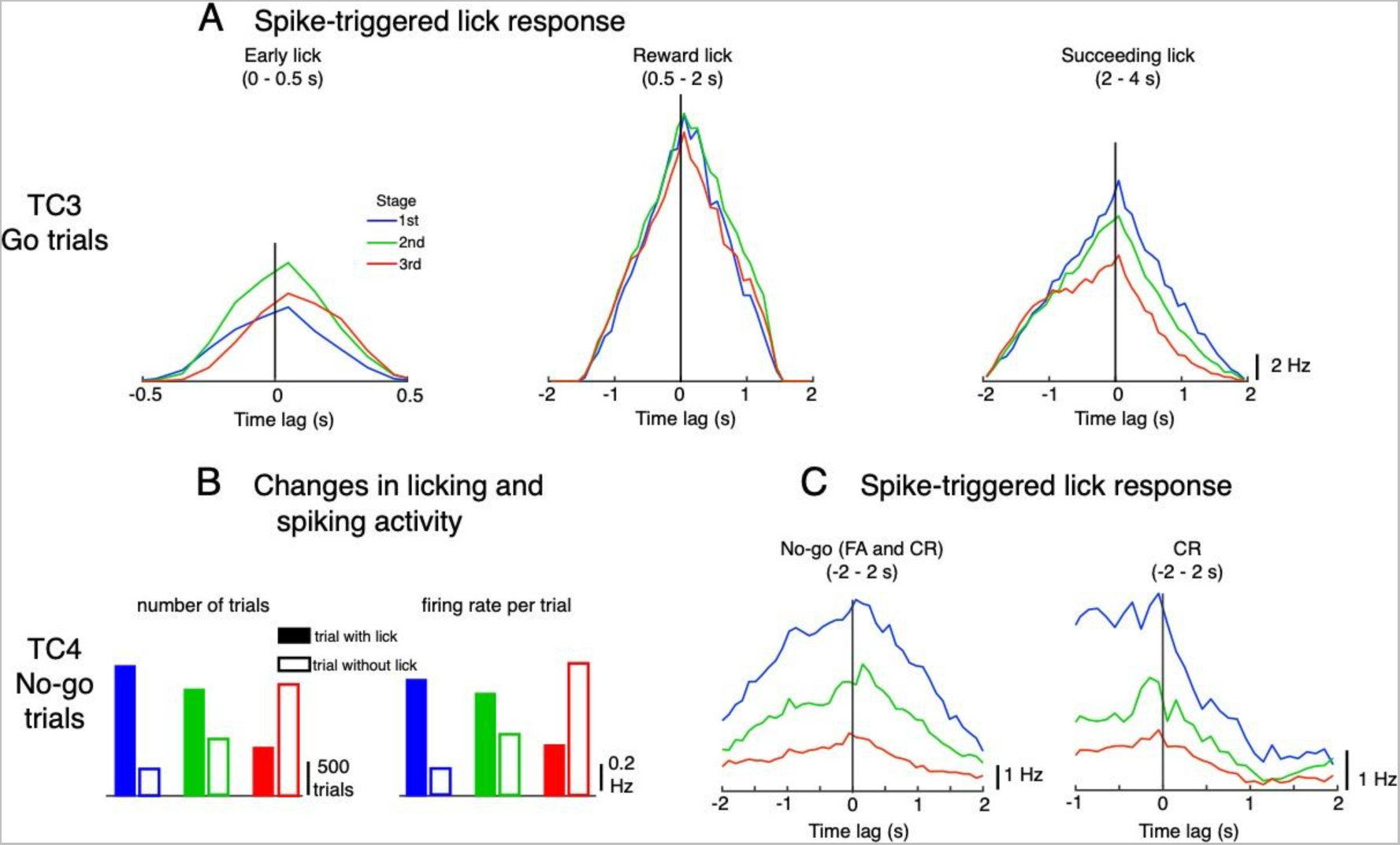
Possible functions of TC3 and TC4. A: spike-triggered lick response for TC3 neurons for Go trials in the three response windows: early lick (0 – 0.5 s), reward lick (0.5 – 2 s) and succeeding lick (2 – 4 s). Lick responses were largest for the reward window with a balanced distribution of negative-positive values, suggesting that TC3 is equally related to sensory feedback and motor control of reward licks. B: change in numbers of trials with (filled bars) and without licks (open bars) during learning, and mean firing rate per trial of TC4 neurons for No-go cues. C: spike-triggered lick responses of TC4 neurons plotted for No-go cues (including FA and CR trials, left) and separately for CR trials (right) in the window −2 to 2 s after cue. Note that according to the experimental design, the subsequent trial was delayed by 1 s from the last lick if the mouse continued to lick. Thus, to compute the spike-triggered lick response of TC4 in −2 to 2s of No-go trials, the transition period of −1 to 0 s, during which there were no licks, was ignored.

**Table S1:**
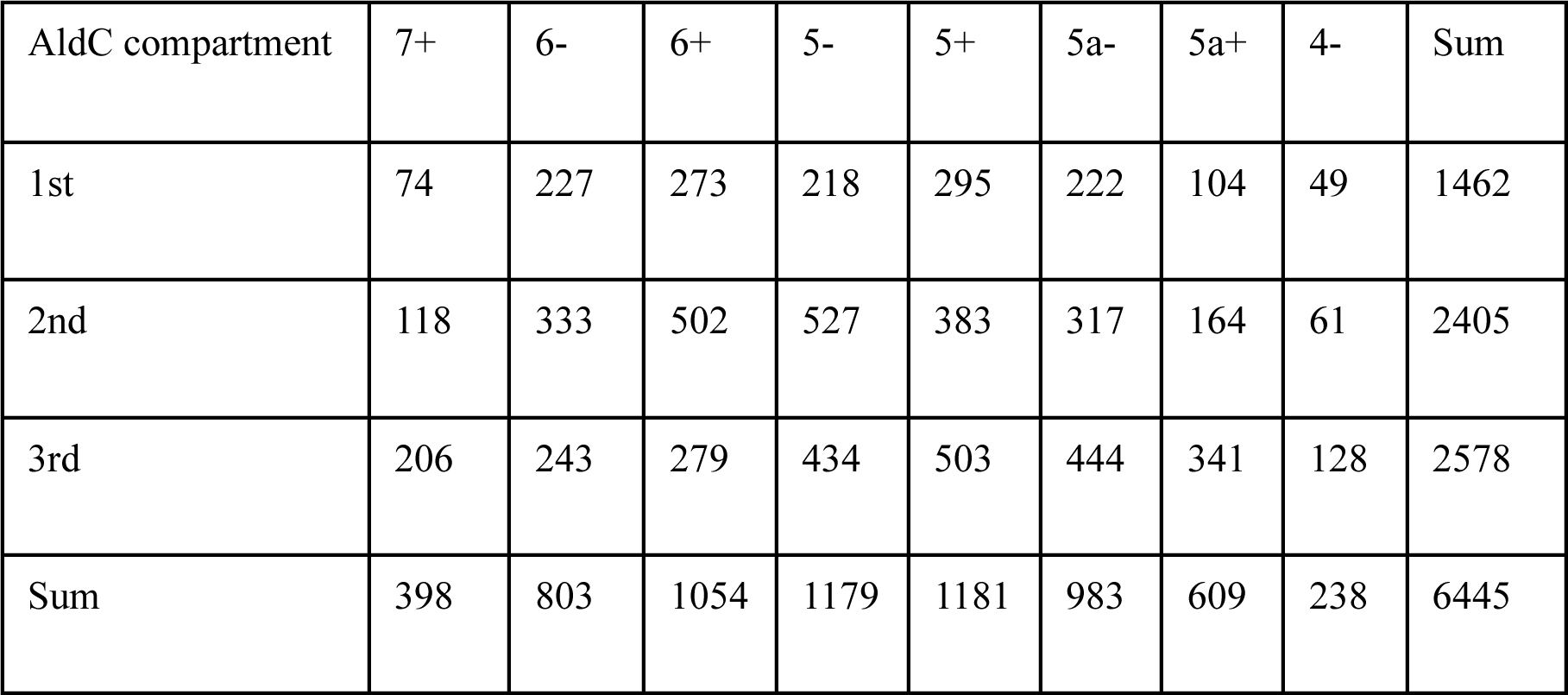
The number of CFs sampled in individual AldC compartments at different learning stages.

**Table S2:**
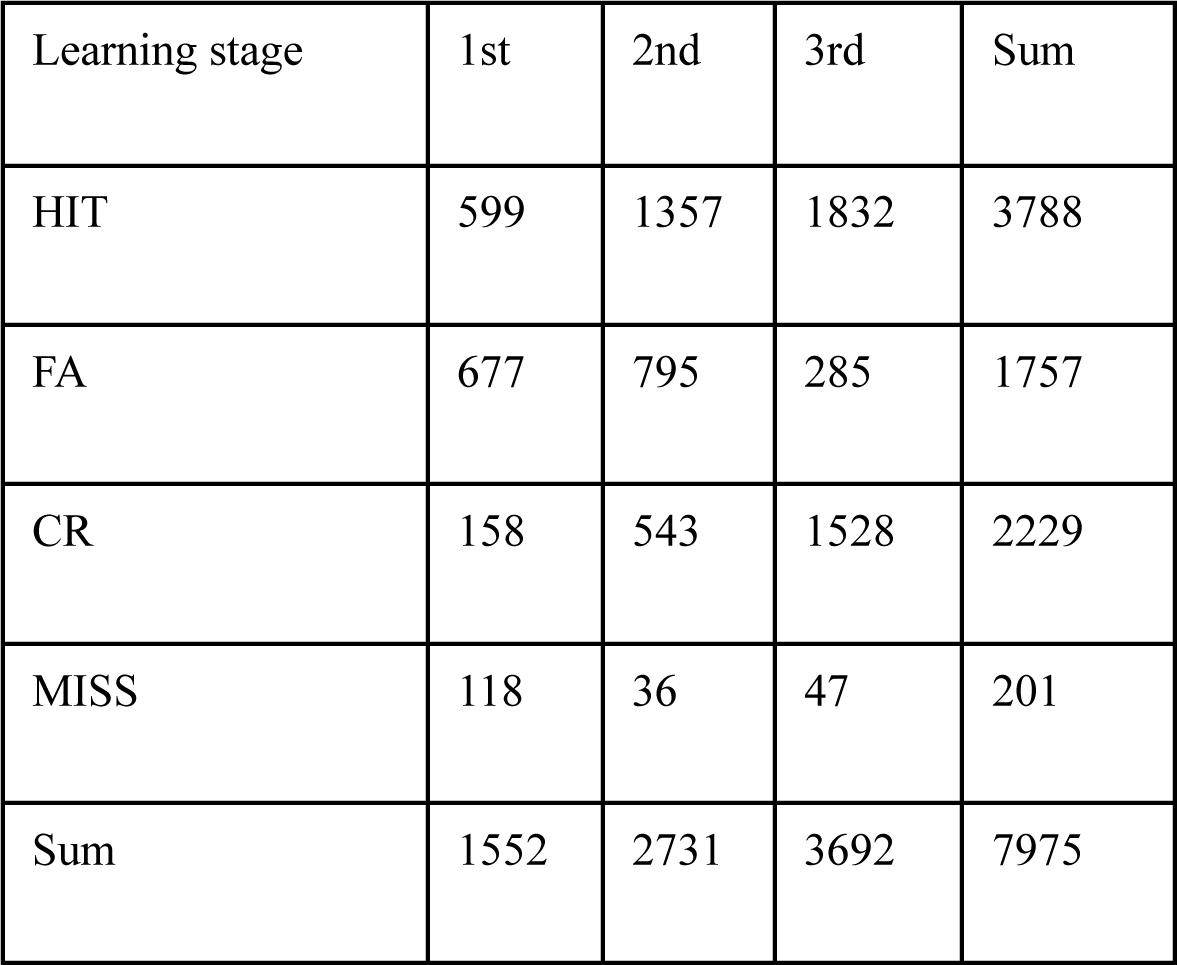
The number of trials for each cue-response condition at different learning stages.

| Learning stage | 1st | 2nd | 3rd | Sum |
| --- | --- | --- | --- | --- |
| HIT | 599 | 1357 | 1832 | 3788 |
| FA | 677 | 795 | 285 | 1757 |
| CR | 158 | 543 | 1528 | 2229 |
| MISS | 118 | 36 | 47 | 201 |
| Sum | 1552 | 2731 | 3692 | 7975 |

**Supplemental Movie M1:** CF firings in 10-ms bins of Ald-C compartment 5+ neurons for HIT trials in two representative sessions of the 1st and 3rd learning stages. Detailed description for the elements can be found in Figure 4C.

**Supplemental Movie M2:** Relationship of TC coefficient, synchrony strength and instantaneous synchrony within individual Ald-C compartment for HIT trials in the 1st and 3rd learning stages of TC1. For each Ald-C compartment (column), the neuron which has highest TC1 coefficients (bottom row) was selected as a reference neuron. Instantaneous synchrony in a single trial (top row) and synchrony strength (second row) were estimated between the reference neuron and other neurons in the same compartment. While synchrony strength was static within session, instantaneous synchrony varies trial-to-trial with strong values observed for HIT trials but not trials of the other cue-response conditions.

